# Boundary conditions for synaptic homeodynamics during the sleep-wake cycle

**DOI:** 10.1101/2024.08.14.607872

**Authors:** Fukuaki L. Kinoshita, Rikuhiro G. Yamada, Koji L. Ode, Hiroki R. Ueda

## Abstract

Understanding synaptic dynamics during the sleep-wake cycle is crucial yet remains controversial. The synaptic homeostasis hypothesis (SHY) suggests synaptic depression during non-rapid eye movement (NREM) sleep, while other studies report synaptic potentiation or synaptic changes during NREM sleep depending on activities in wakefulness. To find boundary conditions between these contradictory observations, we focused on learning rules and firing patterns that contribute to the synaptic dynamics. Using computational models, we found that under Hebbian and spike-timing dependent plasticity (STDP), wake-like firing patterns decrease synaptic weights, while sleep-like patterns strengthen synaptic weights. We refer to this tendency as Wake Inhibition and Sleep Excitation (WISE). Conversely, under Anti-Hebbian and Anti-STDP, synaptic depression during NREM sleep was observed, aligning with the conventional synaptic homeostasis hypothesis. Moreover, synaptic changes depended on firing rate differences between NREM sleep and wakefulness. We provide a unified framework that could explain synaptic homeodynamics under the sleep-wake cycle.

## Introduction

During wakefulness, organisms perceive external worlds through the five senses to learn and take appropriate actions. During sleep, organisms disconnect from the environment to reorganize memory and recover from fatigue. Recent studies have revealed that cortical neurons are responsible for these brain functions underlying learning and memory formation (1) and that the dynamics of the cortical synaptic weights are associated with the sleep-wake cycle (2, 3). A hypothesis known as the synaptic homeostasis hypothesis (SHY) proposed that wakefulness potentiates synapses through learning with the costs of higher energy demand while the sleep state depresses less important synapses to restore synaptic homeostasis (2). However, the dynamics of synaptic weights in the sleep-wake cycle, especially during sleep, remain controversial. Several studies have demonstrated that NREM sleep potentiates synapses, contributing to memory consolidation (4, 5). Furthermore, SHY suggests that synapses strengthened during wakefulness are less susceptible to synaptic depression during NREM sleep (2). In contrast, other studies propose the normalization of neuronal activities, where fast-firing neurons and slow-firing neurons during wakefulness are weakened and strengthened, respectively, during NREM sleep (6). We sought to find the boundary conditions that reconcile these discrepancies and to comprehensively understand the sleep-wake synaptic dynamics.

The heterogeneity of brain states can confound in vivo studies because the slow-wave oscillation (SWO), that is a characteristic firing pattern of NREM sleep, also occurs in wakefulness and sleep states include REM sleep, which has wake-like firing patterns (7, 8). To address this issue, we developed computational models to investigate the direct relationships between synaptic weights and neuronal firing patterns characteristic of each brain state. To account for the diversity of neurons, we prepared various sleep- and wake-like firing patterns based on in vivo experiments. Additionally, we devised a unified function that recapitulates typical synaptic learning rules in the cortex for updating the synaptic weights (9). These settings allowed us to simulate the dynamics of synaptic weights under specific types of spike trains, such as burst firing, based on synaptic learning rules (10). The synaptic learning rules we studied include the Hebbian rule, STDP, and their reverse types (Anti-Hebbian and Anti-STDP). According to the Hebbian rule, a synaptic connection between two neurons strengthens when pre- and post-synaptic neurons fire simultaneously (11). A temporally asymmetric form of Hebbian rule is STDP. The classical STDP describes that synaptic potentiation occurs when pre-synaptic spikes precede post-synaptic spikes within a certain temporal window, while synaptic depression occurs in post-synaptic spikes precede pre-synaptic spikes (12, 13). Reverse types of them (Anti-Hebbian and Anti-STDP) are also observed in the mammalian cortex (14) and play important roles in information processing (15, 16).

Our simulations revealed that synaptic weights become higher in sleep-like synchronized states than in wake-like desynchronized states under Hebbian and classical STDP, assuming the same mean firing rates for both sleep- and wake-like firing patterns. We refer to these dynamics as Wake Inhibition and Sleep Excitation (WISE). In contrast, synaptic depressions during sleep-like firing patterns, which represents SHY, were observed under Anti-Hebbian and Anti-STDP. Moreover, our results suggested that the synaptic dynamics also depend on mean firing rates, providing a unified framework for the synaptic homeodynamics of neural networks during the sleep-wake cycle.

## Results

### WISE under Hebbian and STDP, SHY under Anti-Hebbian and Anti-STDP

To investigate synaptic dynamics in NREM sleep and wakefulness with synaptic learning rules, we used a Ca^2+^-based plasticity model. Graupner et al. proposed that the Ca^2+^-based plasticity model with two thresholds for post-synaptic Ca^2+^ can describe the various types of synaptic learning rules (17). Based on Graupner’s model, we developed a modified Ca^2+^-based plasticity model to represent four different types of learning rules (Hebbian, STDP, Anti-Hebbian, and Anti-STDP) by setting eight parameters (*θ*_*p*_: potentiation threshold, *θ*_*d*_: depression threshold, *γ*_*p*_: potentiation amplitude, *γ*_*d*_: depression amplitude, *τ*_*pre*_: time constant for Ca^2+^ from N-methyl-D-aspartate receptor (NMDAR), *τ*_*post*_: time constant for Ca^2+^ from Voltage-gated Ca^2+^ channel (VGCC), σ: amplitude for noise, and *τ*_*s*_: time constant for synaptic change) (**Fig. 1*A***). Synaptic weights were defined as being linearly related to synaptic efficacy (ρ) as w=w0+ρ(w1-w0), where w0 and w1 are minimum and maximum synaptic weights, respectively. ρ is described by a first order differential equation according to the previous article (*Materials and Methods*) (17). We confirmed that our model could predict the experimentally observed changes of post-synaptic Ca^2+^ and synaptic strength under stimulations with different time lags (**Fig. 1 *B*** and ***C***). We randomly generated more than 1 million parameter sets and selected 1,000 parameter sets that well represent either one of four learning rules (**Fig. 1 *D*** and ***E***, see *Supplementary Materials, Parameter search for synaptic learning rules*). Each learning rule has a clear cluster in the distribution of thresholds and amplitudes (**Fig. 1*F***). The distributions of other parameters and those in other fitting conditions are shown in *Supplementary Materials*, **Fig. S1**. To evaluate the change of synaptic weights during sleep-like and wake-like firing patterns, we assumed one post-neuron connected with ten pre-neuronal synapses and the same mean firing rates both in sleep-like and wake-like patterns (**Fig. 1*G***). Sleep-like and wake-like firing patterns were derived from previous in vivo recordings (6, 18) (*Supplementary Materials, Generation of sleep and wake-like spike patterns* and **Fig. S18**). Then, time changes in synaptic efficacy during the sleep-like and wake-like firing patterns were calculated and compared under synaptic learning rules (*Materials and Methods*). The mean synaptic efficacy became higher in sleep-like states than in wake-like states under Hebbian and STDP, representing WISE (**Fig. 1*H***). The opposite results were observed in Anti-Hebbian and Anti-STDP, representing SHY. Thus, WISE and SHY are observed under the specific types of learning rules, with both states exhibiting equal mean firing rates.

**Figure 1.**
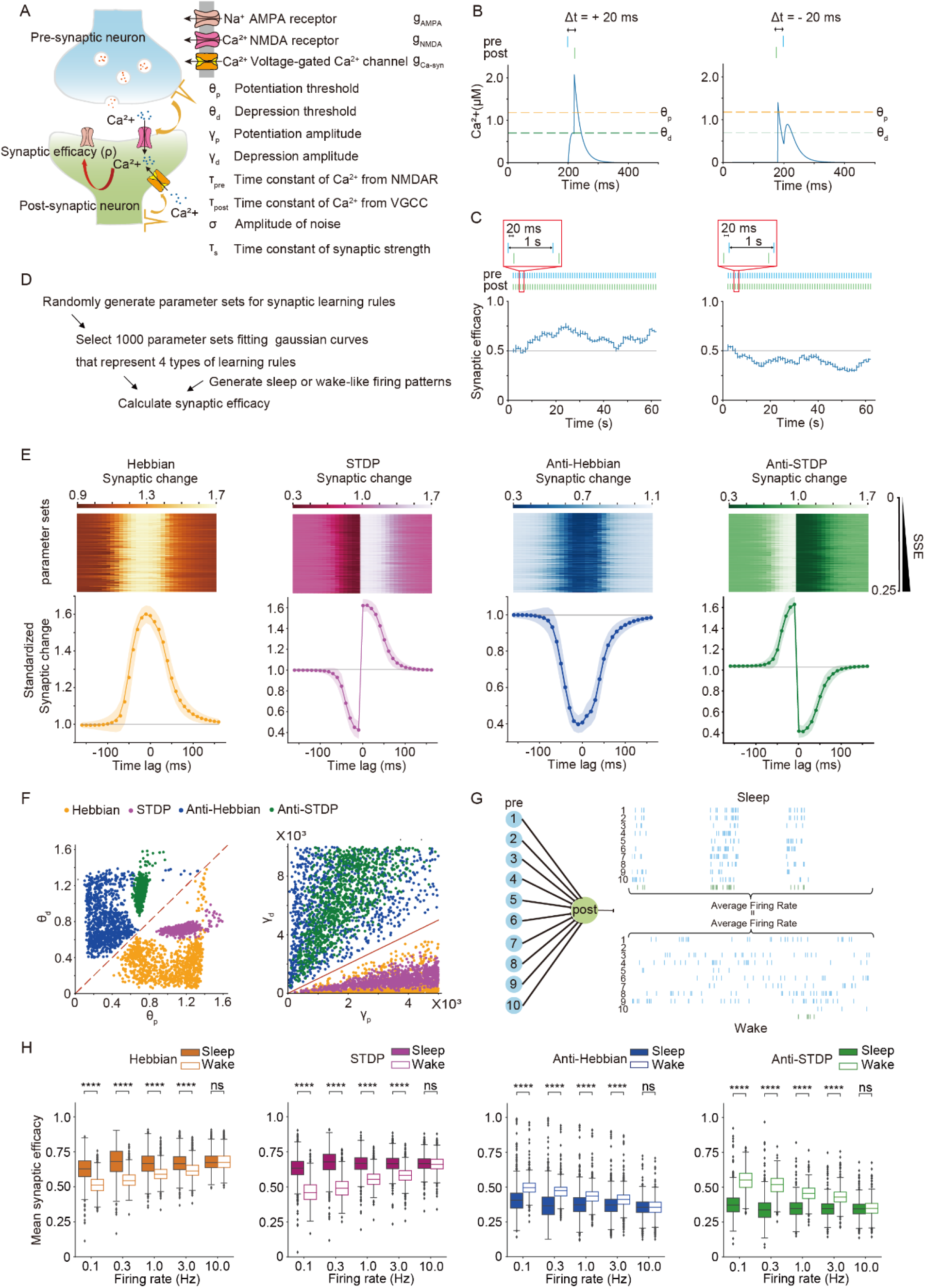
WISE under Hebbian and STDP, SHY under Anti-Hebbian and Anti-STDP assuming the same firing rates during sleep-like and wake-like firing patterns. (A) Schematic illustration of the Ca^2+^-based plasticity model for synaptic learning rules. (B) Calcium transients in a post-neuronal synapse when stimulated with short delays under STDP. (C) The dynamics of synaptic efficacy when stimulated with short delays under STDP. (D) The procedure of a parameter search for synaptic learning rules and calculation of synaptic efficacy in the state of sleep and wake. (E) The standardized synaptic change of the 1000 parameter sets for four types of learning rules represented by Gaussian curves. Each row of the upper panels shows synaptic changes simulated with a parameter set whose sum of squared errors (SSE) between analytical results and Gaussian curves was less than 0.25. The line and shadow in the lower panel indicate the mean and standard deviation, respectively. (F) Distributions of 1000 parameter sets for each learning rule in the axes of thresholds and amplitudes. (G) Schematic illustration for connections and firing patterns of neurons used in the calculation of synaptic efficacy. (H) Box plots for mean synaptic efficacy in sleep-like and wake-like firing patterns by different synaptic learning rules and mean firing rates (*n* = 1000 for each firing rate, *n* represents the number of synaptic learning rules). The whiskers above and below show minimal to maximal values. The box extends from the 25th to the 75th percentile, and the middle line indicates the median. * p < 0.05, ** p < 0.01, *** p < 0.001, **** p < 0.0001, Welch’s t-test was applied.

### Robustness of WISE and SHY

Next, we investigated the robustness of the WISE and SHY in different settings. We first simply modified the previous model with one post-neuron connected by ten pre-neuronal synapses to have 96 pre-neuronal synapses or random connections. This modified model still exhibited WISE under Hebbian and STDP, while it showed SHY under Anti-Hebbian and Anti-STDP (**Fig. 2 *A*** and ***B***). Next, we tested modified parameters of time constants and amplitudes of learning rules to see if WISE and SHY depend on the properties of learning rules. Even in those settings, we found that these trends still held (**Fig. 2*C***, and *Supplementary Materials*, **Fig. S3**). In **Fig. 2 *D*** and ***E***, we generated various sleep-like firing patterns by changing parameters such as the log10(*mean inter-spike interval*) (ISIM), log10(*mean Up-state duration*) (UPM), and log10(*mean Down-state duration*) (DOWNM). We tested ranges of parameters for each targeted mean firing rate by changing either DOWNM (**Fig. 2*D***) or ISIM (**Fig. 2*E***). We found that the mean synaptic efficacy was higher in sleep-like states with most of the parameters in the tested range. Notably, this trend was more apparent at lower firing rates. These results validated that WISE and SHY were robust under various biologically feasible conditions.

**Figure 2.**
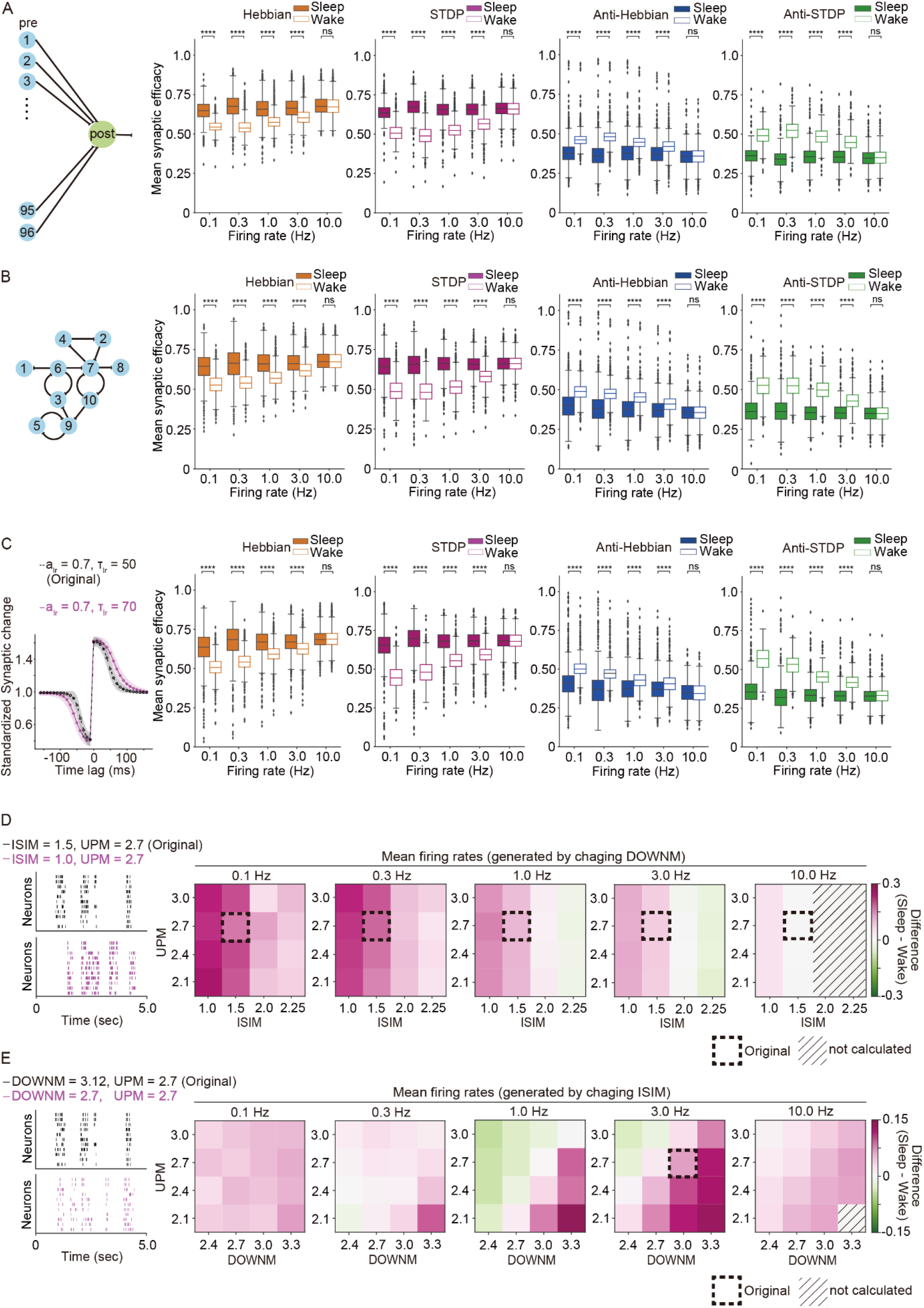
Robustness of WISE and SHY. (A) Schematic illustration of network model consisting of one post-neuron connected with 96 pre-neuronal synapses and box plots for synaptic efficacy. (B) Schematic illustration of neural network model consisting of randomly connected ten neurons and box plots for synaptic efficacy. (C) The original (black dotted line) and modified (magenta solid line) curves for STDP rule and box plots for mean synaptic efficacy. a_Ir_ is the amplitude, and τ_Ir_ is the time constant for Gaussian curves to be fitted by the parameter search. (D) Representative raster plots of the original (black) and modified (magenta) sleep-like firing patterns and the differences in median of mean synaptic efficacies between sleep-like and wake-like firing patterns (*sleep - wake*). The modified mean firing rates of sleep-like firing patterns were generated by changing DOWNM under constant UPM and ISIM (n=1000 for each firing rate). The dotted boxes highlighted the original UPM and ISIM values are 1.5 and 2.7, respectively. (E) Representative raster plots of the original (black) and modified (magenta) sleep-like firing patterns and the differences in median of mean synaptic efficacies between sleep-like and wake-like firing patterns (*sleep - wake*). The modified firing rates of sleep-like patterns were generated by changing ISIM under constant UPM and DOWNM (*n=1000* for each firing rate, *n* represents the number of synaptic learning rules). The dotted boxes highlighted *DOWNM =3*.*0* and *UPM = 2*.*7*, which are the closest to the original values: *DOWNM = 3*.*12* and *UPM = 2*.*7*. (A-C) The mean synaptic efficacy was evaluated in sleep-like and wake-like firing patterns with various mean firing rates assuming different synaptic learning rules (*n = 1000* for each firing rate, *n* represents the number of synaptic learning rules). The whiskers above and below of box plots show minimal and maximal values, respectively. The box extends from the 25th to the 75th percentile and the middle line indicates the median. * p < 0.05, ** p < 0.01, *** p < 0.001, **** p < 0.0001, Welch’s t-test was applied. (D and E) ISIM: log10(*mean ISI*), UPM: log10(*mean Up-state duration*), DOWNM: log10(*mean Down-state duration*)

### WISE and SHY in Hodgkin-Huxley-based network models

We then tested whether WISE and SHY hold in a more realistic setting where the synaptic efficacy can change the firing pattern. To recapitulate the variable firing pattern, we introduced the Hodgkin-Huxley model to network models based on our previous study (19, 20), with a ratio of excitatory and inhibitory neurons of 4:1 (**Fig. 3*A***, see *Supplementary Materials, Hodgkin-Huxley-based network model*). In this study, we considered molecules responsible for generating SWO in three subcellular compartments (post-synaptic, intracellular (cell body), and pre-synaptic compartments) (19, 21). We assumed that NMDAR and VGCC function in post-synapses and cell bodies, respectively, for generating SWO (19). In addition, because pre-synaptic transmission is crucial for generating SWO in the cortex (21), we modeled AMPAR, NMDAR, and GABAR to receive the neural transmission. Parameter searches for SWO and bifurcation analysis for a single neuron were based on previous articles (19). First, we randomly generated parameter sets from a large parameter space within the biophysically feasible range and searched for parameter sets that yield firing patterns of SWO. Then, we searched for parameter sets that bifurcate from wake-like to sleep-like patterns as a network (**Fig. 3*B***, see *Supplementary Materials, Parameter search for SWO and bifurcation analysis in Hodgkin-Huxley-based network models*). Clear desynchronization and synchronization, evaluated by the coefficient of variance (CV) of spike counts per 50 milliseconds (sleep score) (see *Supplementary Materials, Evaluation of synchronization and desynchronization in Hodgkin-Huxley-based network models*), were observed with parameter sets for each subcellular component (**Fig. 3 *C*** and ***D***, and Supplementary Materials, **Fig. S4**). We then selected the parameter sets that showed almost the same mean firing rates in sleep-like and wake-like states and evaluated synaptic efficacy in the sleep-like and wake-like states (*Supplementary Materials*, **Figs. S6** and **S7**). WISE was observed under STDP (**Fig. 3*E***), while SHY was observed under Anti-STDP (**Fig. 3*F***). Similar trends were observed under Hebbian and Anti-Hebbian and in network models with different connections (*Supplementary Materials*, **Figs. S9** and **S10**). These results validated WISE and SHY in a realistic network model.

**Figure 3.**
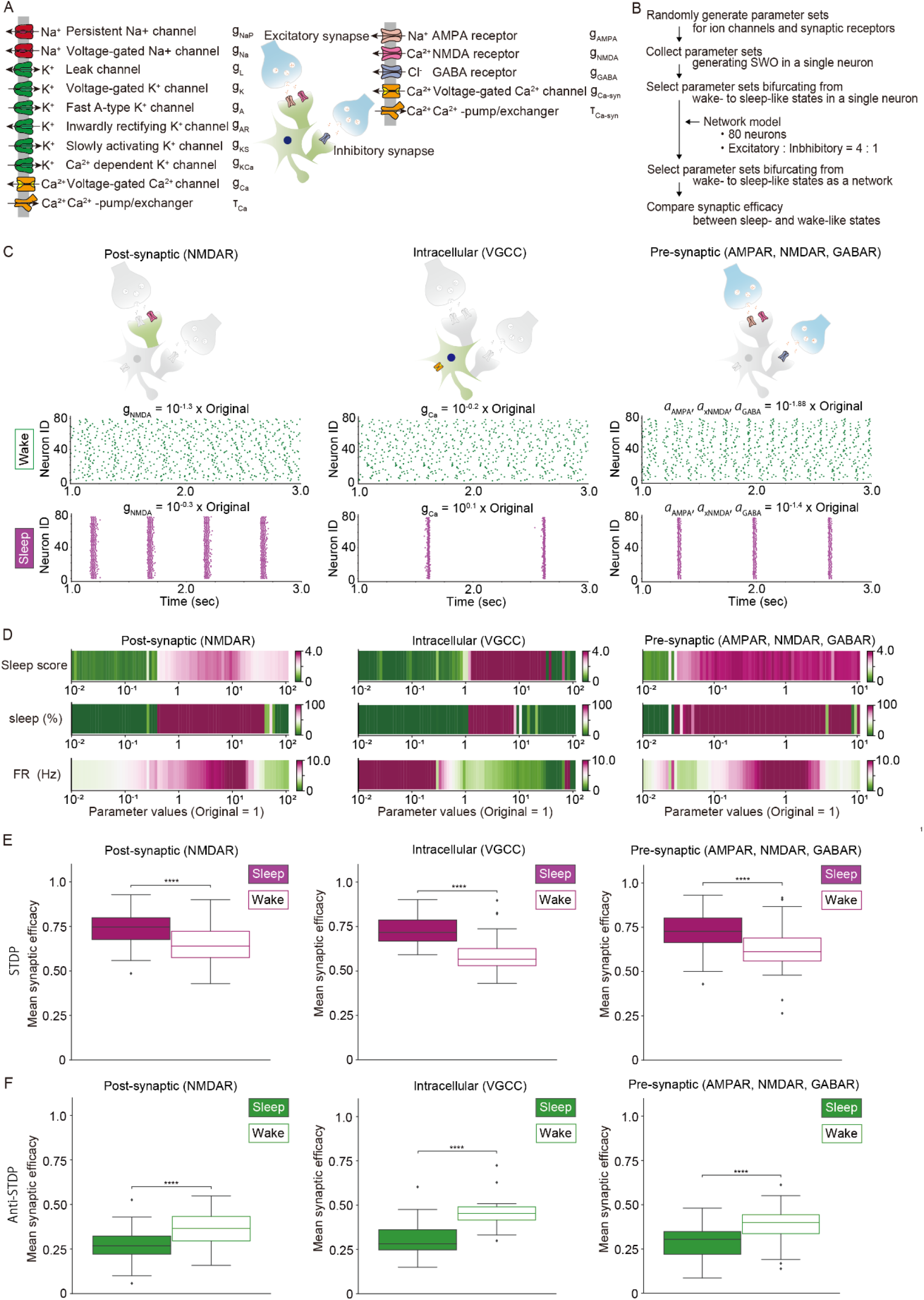
WISE and SHY in Hodgkin-Huxley-based network models. (A) Schematic illustration of a model with excitatory and inhibitory synapses. A Hodgkin-Huxley-based network model was constructed based on the averaged neuron model in a previous study (19). (B) Procedures for collecting parameter sets that bifurcate from wake-like to sleep-like firing patterns and comparing synaptic efficacy between the two states in Hodgkin-Huxley-based network models. (C) Representative sleep-like and wake-like firing patterns for three types of bifurcation models. The parameters used in the simulation were obtained by multiplying the original values defined in **Table S5** by the presented factors. Simulations were conducted for five seconds and presented for three seconds in each model. (D) Changes in the sleep score, percentage of sleep-like waveforms (sleep (%)), and mean firing rate (FR (Hz)) as the conductance of the channel or receptor, or the coefficients of pre-synaptic activations are gradually increased. The range of the conductance was divided into 80 steps for post-synaptic or intracellular bifurcation, while the range of the coefficient was divided into 75 steps for pre-synaptic bifurcation. Simulations were conducted for ten seconds at each conductance or coefficient step. (E and F) Box plots for mean synaptic efficacy during sleep-like and wake-like firing patterns under STDP (E) and Anti-STDP (F) by three types of network models *(n* = 191, 52, and 150 for STDP and n= 121, 36, and 119 for Anti-STDP in post-synaptic, intracellular, and pre-synaptic bifurcation models, respectively. *n* represents the number of parameter sets for the network models). The whiskers above and below of box plots show minimal and maximal values, respectively. The box extends from the 25th to the 75th percentile, and the middle line indicates the median. * p < 0.05, ** p < 0.01, *** p < 0.001,**** p < 0.0001, Student’s t-test was applied.

### WISE under Hebbian and STDP and SHY under Anti-Hebbian and Anti-STDP is compatible with models including sleep-wake dynamics

Next, we incorporated the spontaneous sleep-wake cycle into our network models. Previous phosphoproteomic studies suggested that phosphorylation of several synaptic proteins is associated with sleep needs (22–24). The sleep needs increase during wakefulness and decreases with the onset of sleep. This homeostatic oscillation of sleep needs is referred to as Process S (25). In the present model, we assumed that calcium/calmodulin-dependent protein kinase II (CaMKII) is responsible for the homeostatic oscillation (26). To evaluate the synaptic efficacy across a series of sleep-wake cycles, we assumed that CaMKII changes its states in a use-dependent manner during wakefulness and induces SWO by interacting with channels or receptors that regulate neuronal membrane potentials or enzymes that regulate neurotransmitters (8, 26). This assumption aligns with observations that CaMKII has multiple phosphorylation states and changes its function accordingly (27). We integrated the use-dependent change of CaMKII’s function for activating channels such as NMDAR into the sleep-wake dynamics model (*Materials and Methods*). The initial state of CaMKII, such as pT286/287 CaMKII, has self-activating ability and Ca^2+^-dependent activation (28). This initial state activates the second state of CaMKII, such as pT305/306 CaMKII, or phosphatases, such as calcineurin, which can be regulated during sleep (29). We assumed that the second state directly interacts with molecules that induce SWO from the result of optimizing correlation to Process S for network models (*Supplementary Materials*, **Fig. S11*A***).

In the simulation of a representative model, we observed that pT286/287 CaMKII, represented as ***r*** in **Fig. 4*A***, gradually increased due to Ca^2+^ influx and autoactivation during wakefulness, which was followed by an increase in pT305/306 CaMKII, represented as ***a*** in **Fig. 4*A***. The increased ***a*** then activated NMDAR and induced SWO (**Fig. 4 *C*** and ***D***). The mean synaptic efficacy was higher during sleep-like periods than during wake-like periods under STDP (**Fig. 4 *E*** and ***G***). The opposite results were observed under Anti-STDP (**Fig. 4 *F*** and ***H***). Statistical significances were observed in the analysis with multiple parameter sets (*Supplementary Materials*, **Fig. S11 *B*** and ***C***). The results of model with other bifurcation mechanisms or systems also showed the same trends *(Supplementary Materials*, **Figs. S12**-**S14**). These results confirmed that WISE under STDP and SHY under Anti-STDP are compatible with network models that have sleep-wake dynamics.

**Figure 4.**
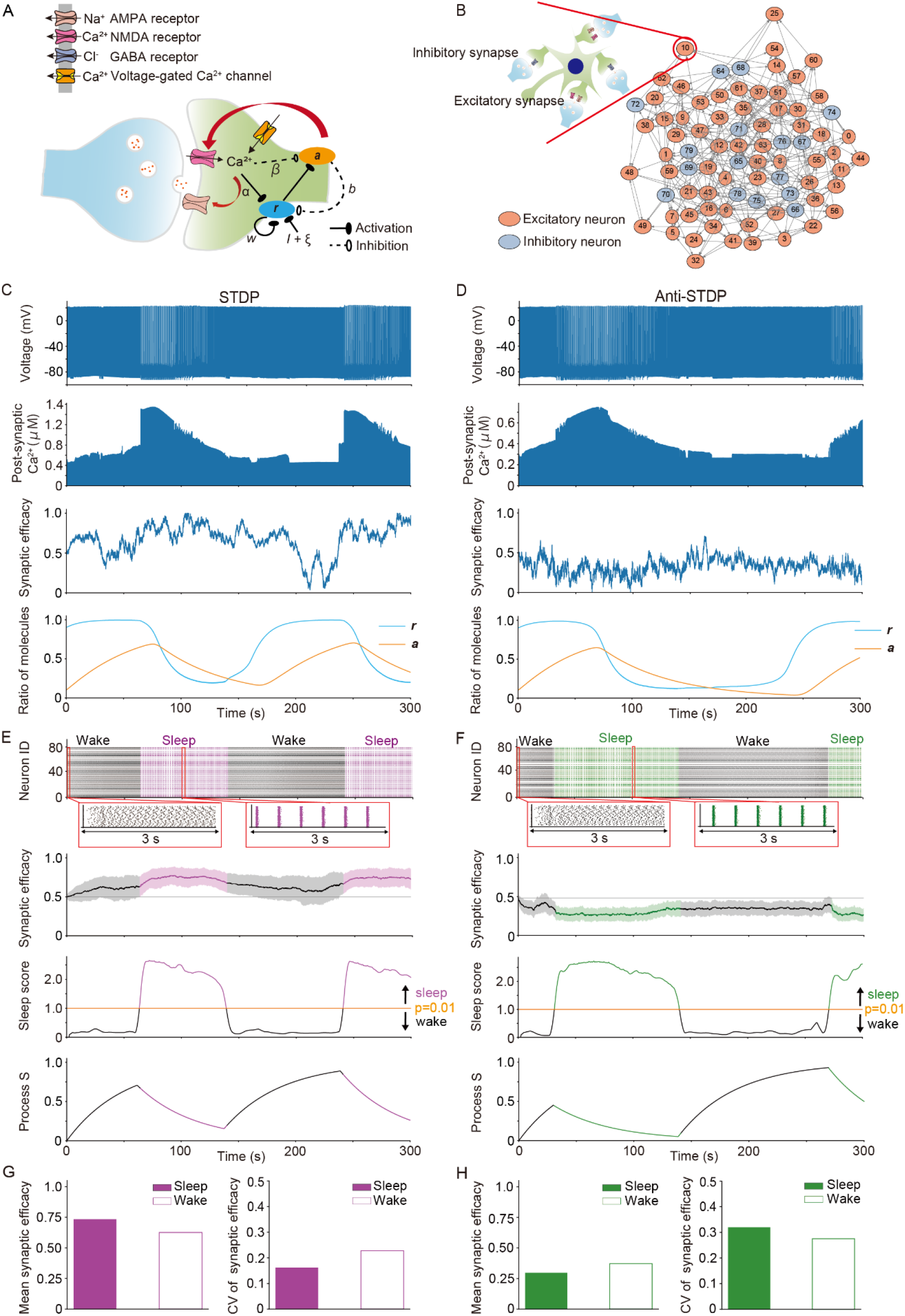
WISE under Hebbian and STDP, and SHY under Anti-Hebbian and Anti-STDP is compatible with models including sleep-wake dynamics. (A) Schematic illustration of the model for sleep-wake dynamics in excitatory synapses. The initial state of CaMKII such as pT286/287 CaMKII is represented by ***r***. The initial state activates ***a*** which corresponds to the second state of CaMKII such as pT305/306 CaMKII or phosphatases such as calcineurin. *α, β, w*, and *b* corresponds to variables defined in the equations in the *Materials and Methods*. Synaptic efficacies were calculated in Hodgkin-Huxley-based network models bifurcated by the post-synaptic mechanism with sleep-wake dynamics under synaptic learning rules. The conductance of NMDAR was updated by ***a*** and the simulations were optimized by Pearson’s correlation coefficients between process S and ***r***. The simulations were conducted for 500 seconds. The parameter set for channel or receptor conduces, learning rule and sleep-wake dynamics and initial values for variables in a representative model are shown in **Tables S5-S8**. (B) Schematic illustration of the network model used in simulations. The network model has 80 neurons with the E:1 ratio of 4:1. (C and D) Time changes of post-synaptic membrane potential and Ca^2+^ concentration, synaptic efficacy, and the ratio of two phosphorylated states of kinases (***r*** and ***a***) of a synapse in representative network models under STDP (C) and Anti-STDP (D). The results of 0-300 seconds are shown. (E and F) Raster plots and time changes of mean synaptic efficacy, sleep score, and process S in representative network models under STDP (E) and Anti-STDP (F). The shadow in time changes in mean synaptic efficacy represents SD. The network was considered to be in the state of sleep or wake if the sleep score was above or below the threshold, respectively (the threshold is the value of sleep score where p = 0.01; see *Supplementary Materials, Evaluation of synchronization and desynchronization in Hodgkin-Huxley-based network models*). The results of 0-300 seconds are shown. (G and H) Mean and coefficient of variance (CV) of synaptic efficacy during the periods of sleep-like and wake-like states in representative network models under STDP (E) and Anti-STDP (F).

### Synaptic changes depend on firing rates assuming higher firing rates during wake-like states

In the previous sections, we compared synaptic efficacies in sleep-like and wake-like states by assuming the same mean firing rates in both states. While this assumption is feasible in brain regions such as the visual cortex, where firing rates are almost constant between states (30), regions such as the somatosensory cortex showed higher firing rates during wakefulness (31). To evaluate the synaptic dynamics in the higher firing rates during wake-like states, we assumed that mean firing rates in Up states of sleep-like patterns are equal to those in wake-like patterns and found that WISE was observed at lower mean firing rates while SHY was observed at higher mean firing rates under Hebbian and STDP (**Fig. 5*A***). Thus, synaptic efficacies change to different directions depending on mean firing rates assuming higher firing rates during wake-like states.

**Figure 5.**
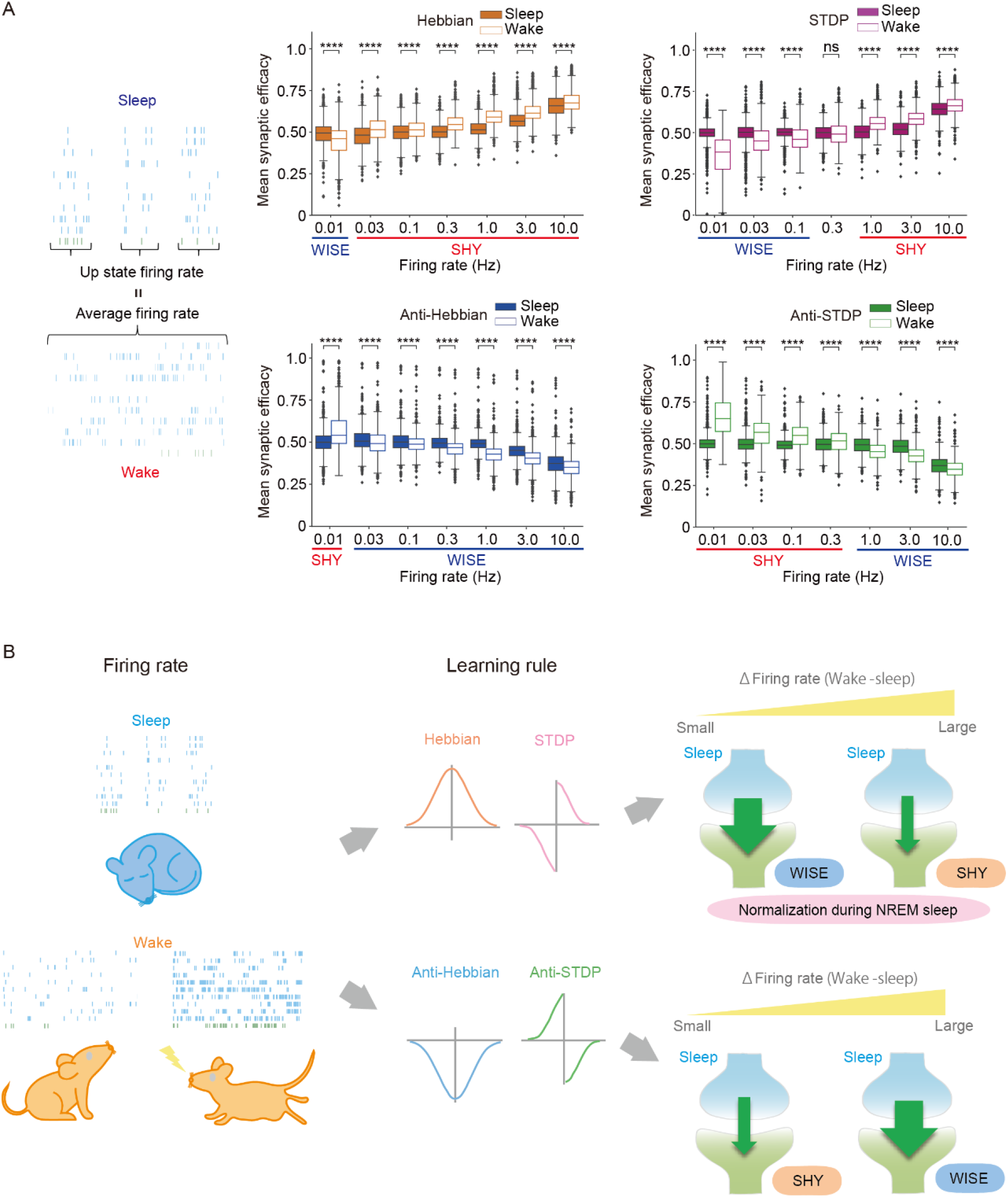
Synaptic changes depend on firing rates assuming higher firing rates during wakefulness. (A) A Schematic illustration for sleep-like and wake-like spike patterns and box plots for mean synaptic efficacy in sleep-like and wake-like firing patterns for different learning rules and mean firing rates (*n* = 1000 for each firing rate, *n* represents the number of synaptic learning rules). The mean firing rates in wake-like states are equal to the mean firing rates in Up states of sleep-like states. The sleep-like firing patterns were generated by sampling from the lognormal distributions with log10(*mean Up-state duration*) = 2.7 and log10(*mean Down-state duration*) = 3.0. SD was calculated according to the linear regression analysis based on in vivo data (see *Supplementary Materials*, **Fig. S18**). WISE and SHY dynamics are highlighted under the x-axis according to the change in the direction of synaptic efficacy. The whiskers above and below show minimal to maximal values. The box extends from the 25th to the 75th percentile and the middle line indicates the median. * p < 0.05, ** p < 0.01, *** p < 0.001, **** p < 0.0001, Welch’s t-test was applied. (B) A Graphical abstract of a unified framework for synaptic dynamics during the sleep-wake cycle. SHY: Synaptic homeostasis hypothesis, WISE: Wake Inhibition Sleep Excitation

## Discussion

In this study, we investigated how synaptic dynamics interact with firing patterns and learning rules. Under Hebbian and STDP, wake-like firing patterns inhibits the synaptic connections, hence weakening the synaptic efficacy, while sleep-like firing patterns excite synaptic connections, hence strengthen the synaptic efficacy. We referred to this tendency as WISE. In contrast, under Anti-Hebbian and Anti-STDP, wake-like and sleep-like patterns tend to strengthen and weaken synaptic efficacies, respectively, which aligns with SHY. When we set the firing rate of the Up state of sleep-like phasic firing patterns equal to the firing rate of wake-like tonic firing patterns, the resulting higher firing rate of wake-like firing pattern tends to strengthen synapses. This indicates that firing rate is the dominant factor in determining the direction of synaptic changes. These findings delineate the boundary conditions of synaptic dynamics during the sleep-wake cycle. We also demonstrated that these boundary conditions are stable under various conditions by using two types of models with numerous parameters derived from biological knowledge.

### Boundary conditions in synaptic homeodynamics

From the perspective of homeostasis, SHY proposes that wakefulness strengthens synapses to learn about the environment, while NREM sleep weakens less important synapses to reduce energy consumption (2). Although studies, including anatomical and electrophysiological research, support SHY (22, 32, 33), several studies have reported contradictory results (4–6, 34).

Our study indicated that the direction of synaptic changes during sleep-like firing patterns depends on synaptic learning rules and firing rates of local networks. Assuming the same mean firing rates in sleep-like and wake-like patterns, SHY is observed under Anti-Hebbian and Anti-STDP (**Figs. 1*H*, 2 *A*-*C*, 3 *E*** and ***F*, and 4 *G*** and ***H***). Additionally, we observed that higher mean firing rates results in smaller differences in synaptic changes between sleep-like and wake-like states (**Figs. 1*H*** and **2 *A-C***, and *Supplementary Materials* **Fig. S9**). This finding suggests that neurons with higher firing rates during wakefulness are less susceptible to synaptic depression during sleep, consistent with SHY (2). In contrast, we observed WISE under Hebbian and STDP (**Figs. 1*H*, 2 *A-C*, 3 *E*** and ***F*, and 4 *G*** and ***H***). This observation indicated that the maximum firing rates during sleep, particularly in the Up state of SWO, are higher when the mean firing rates are equal in both sleep and wakefulness. The high maximum firing rates in sleep enhance synaptic learning rules, that is, Hebbian and STDP strengthen synaptic weights while Anti-Hebbian and Anti-STDP weaken synaptic weights. In vivo experiments also reported that the higher firing rates or shorter inter-spike interval (ISI) in the Up state of SWO (6, 7, 18) (*Supplementary Materials*, **Fig. S18**). Higher maximum firing rates lead to greater Ca^2+^ influx and larger changes in synaptic weights during SWO. Additionally, synchronization and hyperpolarized Down states that promote Ca^2+^ influx in the subsequent Up state (35) likely contribute to elevated post-synaptic Ca^2+^ during SWO.

We propose that synaptic changes depend on the differences in firing rates between NREM sleep and wakefulness. SHY may occur under STDP when mean firing rates during wakefulness are higher than during NREM sleep (**Fig. 5*A***). A simulation showed that synaptic efficacies of neurons which are stimulated during wake-like states change according to SHY (see *Supplementary Materials, Calculation of synaptic efficacy under synaptic learning rules in Hodgkin–Huxley-based network models including stimulation during the wakefulness*, and **Fig. S15**). Similarly, previous studies have demonstrated that exposure to novel stimuli or enforced wakefulness, conditions expected to increase sleep pressure, result in synaptic downscaling during sleep (36, 37). Conversely, quiet conditions, anticipated to yield lower sleep pressure during wakefulness, have strengthened synapses during sleep (36). These observations align with our proposal because exposure to novel environments or higher activity increases firing rates during wakefulness, leading to SHY. In contrast, the quiet wake causes only limited differences in firing rate between NREM sleep and wakefulness, leading to WISE (**Fig. 5*A***). These different responses to the neuronal activities during wakefulness support the idea that NREM sleep normalizes the neuronal activities that are skewed during the wakefulness, as presented by Watson et al (6).

Noteworthy, WISE predicts lower post-synaptic Ca^2+^ concentration during wake-like desynchronized firing under Hebbian and STDP. This prediction aligns with the observation that calcineurin, an LTD-related molecule likely to be activated by lower Ca^2+^ concentration during wakefulness, plays a role in excitatory post-neuronal synapses for generating SWO in the following NREM sleep (38–40). Another prediction of WISE is synaptic connectivity homeostasis. When synaptic transmission is inhibited, the resulting SWO may strengthen synapses through WISE, compensating for the inhibited transmission. The connectivity homeostasis is also anticipated in the synaptic dynamics during hibernation. Decreases in firing rates and synaptic connections due to low temperatures during hibernation are associated with increases in SWO during NREM sleep and restoration of synaptic connections after hibernation (41, 42), that is contrary to SHY. Since the lower the firing rates, the greater the synaptic potentiation during sleep in our results (**Fig. 1*H*** and *Supplementary Materials*, **Fig. S9**), WISE may explain synaptic dynamics during the hibernation. Likewise, in depressive disorder, which is characterized by reduced waking activity and dysfunction of AMPAR in frontal cortex (43), synaptic increase during NREM sleep may occur.

In conclusion, our study provides a unified framework for the synaptic dynamics during the sleep-wake cycle (**Fig. 5*B***). It suggests that SHY, WISE, and the normalization during NREM sleep coexist but occur depending on synaptic learning rules and neuronal activities of networks. Although further studies are needed to investigate relationships between synaptic learning rules, neuronal activities, and synaptic weights, the framework we presented here lays a foundation for future research.

### Synaptic dynamics and brain functions

Our study provides several implications regarding the relationship between synaptic dynamics and brain functions during the sleep-wake cycle. WISE supports the notion that synaptic potentiation in NREM sleep contribute to memory consolidation (4, 34), which can be promoted by STDP (44, 45). On the other hand, we revealed that wake-like desynchronized states can lead to synaptic depression under Hebbian and STDP, especially at lower firing rates (**Figs. 1*H*, 2 *A-C*, 3 *E*** and ***F***, and **4 *G*** and ***H***). One implication of this inhibition during desynchronized states is the enhancement of the signal to noise ratio (SNR) (46, 47). Stimulated neurons likely fire at higher rates, and their synaptic efficacies become larger than those of other neurons in the background desynchronized states. Additionally, neurotransmitter release during wakefulness possibly modulates learning rules to control SNR (48). Indeed, it has been reported that cholinergic projections from the nucleus basalis to the cortex are responsible for synaptic inhibition (47).

We also found that the variability of synaptic efficacy is higher during wakefulness than in NREM sleep under Hebbian and STDP (**Fig. 4*G***, and *Supplementary Materials*, **Figs. S12**-**S14** and **S16**). Several studies suggest that the variability of synaptic efficacies leads to the variability of network activity and reflects probabilistic inference for external worlds and decision-making (49–51). In this context, our results suggest that awake states, with higher variability of synapses, are advantageous for exploration and behavioral selection. Additionally, the decline in synaptic variability during NREM sleep under Hebbian and STDP supports the idea of normalization of firing rates, whose distribution is skewed during wakefulness (6).

### Limitations

The present study has some limitations. Our models are composed of uniform neuronal populations and synaptic learning rules, which can be regulated by sleep promoting kinases. However, different types of neurons are interconnected in the cortex, and their firing rates and synaptic learning rules vary by region and condition (9, 48, 52, 53). Further studies on synaptic plasticity across a wide variety of neurons, regions, and conditions will provide a more detailed understanding of the relationship between synaptic learning rules and synaptic plasticity.

In this study, we assumed that the activation of sleep-promoting kinases and the accumulation of phosphorylation during wakefulness induce SWO. We presented three bifurcation models for the generation of SWO based on the changes in firing patterns (**Figs. 3** and **4**). Since the molecular mechanism of SWO is still under vigorous investigations, these bifurcation models will need to be updated by future experiments. Additionally, the model for the sleep-wake dynamics has some ambiguities (**Fig. 4**). The coefficient “*b*” in the equation (*Eq*. 14, *Materials and Methods*) can represent not only the second phosphorylated states of CaMKII such as pCaMKII (T305/T306), but also phosphatases such as calcineurin (38). Moreover, other kinases such as salt-inducible kinase 3 and extracellular signal-regulated kinase can also correspond to components in the model (54, 55). Further studies to elucidate the interactions between these sleep-promoting kinases and synaptic proteins related to membrane potentials or synaptic strength will contribute to a more accurate understanding of the mechanism for SWO and sleep-wake dynamics.

## Materials and Methods

The details of the materials and methods are supplied in the *Supplementary Materials, Supporting Information Text*.

### Modeling Synaptic Learning Rules

Synaptic efficacy variable ρ is described by a first order differential equation according to the previous article (17).

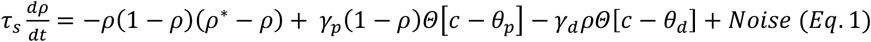

(*Θ* denotes the Heaviside function: *Θ*[*c*−*θ*] = 0 for *c* < *θ* and *Θ*[*c*−*θ*] = 1 for *c* ≥ *θ*).

*c* is the post-synaptic Ca^2+^ concentration and expressed as the sum of Ca^2+^ influx from NMDAR (*C*_*pre*_) and VGCC (*C*_*post*_).

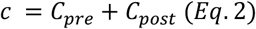

*C*_*pre*_ follows a second order differential equation of NMDAR activation by pre-neuronal firing. We modified the NMDAR kinetics and didn’t consider Mg^2+^ block against NMDAR for the purpose of describing the four different types of learning rules.

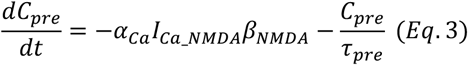

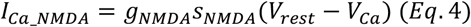

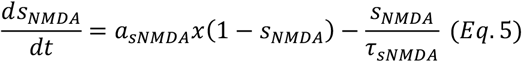

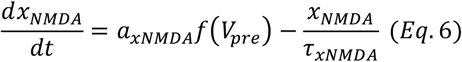

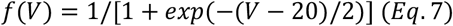

C_post_ is described by a first order differential equation of VGCC

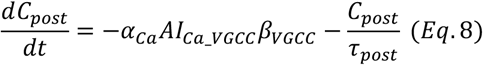

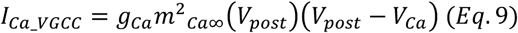

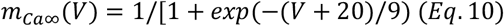

Equations (*Eq*. 3-10) were based on the previous study (1). Fixed parameters are shown in the Table S1. Scaling parameters *β*_*NMDA*_ *and β*_*VGCC*_ in Equations (*Eq*. 3 and 8) were estimated so that the mean amplitude of *C*_*pre*_ = 0.7 μM and the mean amplitude of *C*_*post*_ =1.4 μM according to the results of the previous studies (56). The equation for noise and the comparison of simulation and analytical results are in *Supplementary Materials, Comparison of simulation and analytical results for synaptic learning rules* and **Fig. S17**.

### Comparison of synaptic efficacy during sleep-like and wake-like firing patterns in simple model

The generated sleep-like or wake-like spike trains were converted to the data for membrane potential (see *Supplementary Materials, Conversion from spike patterns to voltage waveforms*). Time changes in synaptic efficacy were calculated by assigning the time-series data for sleep-like and wake-like membrane potentials and parameter sets for synaptic learning rules to the equations (*Eq*. 1-10). Time changes in synaptic efficacy was calculated for 6, 18, and 36 minutes with mean firing rates of pre-synaptic neurons being 0.1-10.0 Hz, 0.03 Hz and 0.01 Hz, respectively. Mean and CV of synaptic efficacy for the last two minutes were calculated by every time step and compared between sleep-like and wake-like firing patterns as an average over time. In random connections of **Fig. 2*B***, each neuron was randomly connected to other neurons with a probability of 12 % and the synaptic efficacy was calculated based on their spike trains.

### Comparison of synaptic efficacy between sleep-like and wake-like firing patterns in Hodgkin-Huxley-based network models

A representative synaptic learning rule was obtained in each network model (see *Supplementary Materials, Parameter search for synaptic learning rules in Hodgkin-Huxley-based network models*). Time changes in synaptic efficacy were calculated for 60 seconds in each of wake-like and sleep-like patterns under specific learning rules in **Fig. 3**. Scaling parameters (*β*_*NMDA*_and *β*_*VGCC*_) and normalized thresholds for learning rules (*θ*_*p*_ and *θ*_*d*_) were calculated by preliminary simulations (see *Supplementary Materials, Obtaining values for scaling parameters and normalization of thresholds for synaptic learning rules*). The mean and CV of synaptic efficacy of excitatory neurons were calculated by every time step except for the initial 10 seconds and compared between sleep-like and wake-like patterns as an average over time. In this process, the network models with large differences of mean firing rates between sleep-like and wake-like patterns (> 2.0 Hz) were excluded from the statistical analysis. Because the AMPAR activity in synapses reflects synaptic strength (57), the conductance of AMPAR was updated by synaptic efficacy (*ρ*).

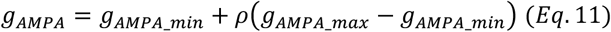

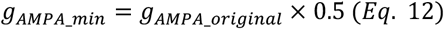

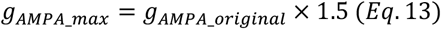

### Calculation of synaptic efficacy under synaptic learning rules in Hodgkin-Huxley-based network models with sleep-wake dynamics

The equations of the two variable model for sleep-wake dynamics are based on the previous study (58).

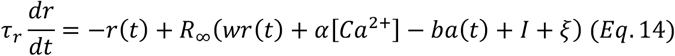

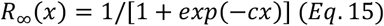

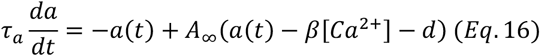

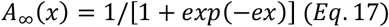

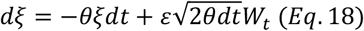

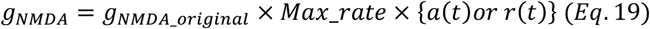

We constructed the network of 80 neurons as in **Fig. 3** including synaptic learning rules and sleep-wake dynamics and calculated the time change of ***r*** and ***a***. Network models analyzed in **Fig. 3 *B-F*** and with continuous transitions from wake-like patterns to sleep-like patterns were selected. For the model of sleep-wake dynamics bifurcated by the pre-synaptic mechanism, we assumed that the pre-synaptic Ca^2+^ ([*Ca*^2+^ _*pre*_]) is mediated by VGCC in pre-neuronal synapses and it was expressed by the following equations.

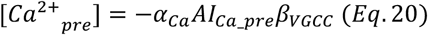

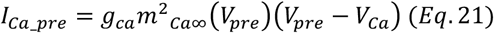

[Ca^2+^] in the equations (*Eq*. 14 and 16) was updated by Ca^2+^ in post-neuronal synapses, cell body or pre-neuronal synapses in the bifurcation model of the post-synaptic, intracellular or pre-synaptic mechanism, respectively. Sleep scores were calculated for 5-seconds’ windows and moving averages of them were computed on 10 consecutive values, which were classified into sleep-like and wake-like periods if sleep score was more than or less than the threshold, respectively (threshold is the value of sleep score when p < 0.01, see *Supplementary Materials, Evaluation of synchronization and desynchronization in Hodgkin-Huxley-based network models*).

Then, we calculated Process S as following equations proposed by a previous study (59).

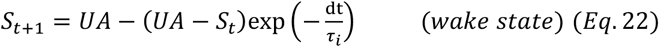

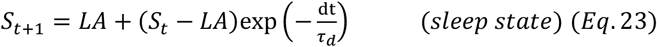

We selected the models in which periods of sleep-like and wake-like states were close to each other (0.7 <= total sleep time/total wake time < 1.3) and Pearson’s correlation coefficient between ***r*** or ***a*** and Process S was the largest. Scaling parameters (*β*_*NMDA*_ and *β*_*VGCC*_) and normalized thresholds for learning rules (*θ*_*p*_ and *θ*_*d*_) were calculated by preliminary simulations (see *Supplementary Materials, SI Materials and Methods, Obtaining values for scaling parameters and normalization of thresholds for synaptic learning rules*). The simulations were conducted for 300 or 500 seconds. The mean and CV of synaptic efficacy of all neurons were calculated by every time step during the periods of sleep-like and wake-like states and compared as an average over time.

## Acknowledgments

We thank all the laboratory members at RIKEN Center for Biosystems Dynamics Research and the University of Tokyo. We thank M. Graupner for guidance of codes. We thank B. O. Watson and D. Levenstein for guidance of in vivo datasets. We thank MN Ballester Roig for reviewing manuscripts. We thank G. Buzsáki, V. V. Vyazovskiy, C. Cirelli, G. H. Diering, P. Meerlo, P. Franken, M. G. Frank, A. Adamantidis, H. C. Heller, M. Schmidt and A. Loudon for helpful discussions. This study was supported by JST ERATO (grant number JPMJER2001, to H.R.U.); the Science and Technology Platform Program for Advanced Biological Medicine (AMED/MEXT to H.R.U.); a Grant-in-aid for scientific research (S) (to H.R.U., grant number JP18H05270); MEXT Quantum Leap Flagship Program (MEXT QLEAP) (to H.R.U., grant number JPMXS0120330644); a Grant-in-Aid from the Human Frontier Science Program (to H.R.U.); RIKEN Junior Research Associate Program (to F.L.K.).

## Author Contributions

F.L.K., R.G.Y., and H.R.U. designed the study; F.L.K. performed the most of simulation study; F.L.K., R.G.Y., and K.L.O. validate the results; F.L.K., R.G.Y., K.L.O., and H.R.U wrote the manuscript.

## Competing Interest Statement

The authors declare that they have no competing interests.

## Notes

### Competing Interest Statement

The authors have declared no competing interest.

### Summary of Updates

Abstract and Figure legends are revised.

## References

1. K. H. Jawabri, S. Sharma, Physiology, Cerebral Cortex Functions (StatPearls Publishing, 2023).

2. G. Tononi, C. Cirelli, Sleep and the Price of Plasticity: From Synaptic and Cellular Homeostasis to Memory Consolidation and Integration. Neuron 81, 12–34 (2014).

3. J. Seibt, M. G. Frank, Primed to Sleep: The Dynamics of Synaptic Plasticity Across Brain States. Front. Syst. Neurosci. 13, 2 (2019).

4. S. Chauvette, J. Seigneur, I. Timofeev, Sleep oscillations in the thalamocortical system induce long-term neuronal plasticity. Neuron 75, 1105–1113 (2012).

5. I. Timofeev, S. Chauvette, Sleep slow oscillation and plasticity. Curr. Opin. Neurobiol. 44, 116–126 (2017).

6. B. O. Watson, D. Levenstein, J. P. Greene, J. N. Gelinas, G. Buzsáki, Network Homeostasis and State Dynamics of Neocortical Sleep. Neuron 90, 839–852 (2016).

7. M. Steriade, I. Timofeev, F. Grenier, Natural waking and sleep states: a view from inside neocortical neurons. J. Neurophysiol. 85, 1969–1985 (2001).

8. V. V. Vyazovskiy, et al., Local sleep in awake rats. Nature 472, 443–447 (2011).

9. D. E. Feldman, The spike-timing dependence of plasticity. Neuron 75, 556–571 (2012).

10. P. J. Sjöström, G. G. Turrigiano, S. B. Nelson, Rate, timing, and cooperativity jointly determine cortical synaptic plasticity. Neuron 32, 1149–1164 (2001).

11. D. O. Hebb, The Organization of Behavior: A Neuropsychological Theory (Psychology Press, 2005).

12. G. Chechik, Spike-timing-dependent plasticity and relevant mutual information maximization. Neural Comput. 15, 1481–1510 (2003).

13. Y. Dan, M.-M. Poo, Spike timing-dependent plasticity of neural circuits. Neuron 44, 23–30 (2004).

14. G. Koch, V. Ponzo, F. Di Lorenzo, C. Caltagirone, D. Veniero, Hebbian and anti-Hebbian spike-timing-dependent plasticity of human cortico-cortical connections. J. Neurosci. 33, 9725–9733 (2013).

15. P. D. Roberts, C. V. Portfors, Design principles of sensory processing in cerebellum-like structures. Early stage processing of electrosensory and auditory objects. Biol. Cybern. 98, 491–507 (2008).

16. P. D. Roberts, T. K. Leen, Anti-hebbian spike-timing-dependent plasticity and adaptive sensory processing. Front. Comput. Neurosci. 4, 156 (2010).

17. M. Graupner, N. Brunel, Calcium-based plasticity model explains sensitivity of synaptic changes to spike pattern, rate, and dendritic location. Proc. Natl. Acad. Sci. U. S. A. 109, 3991–3996 (2012).

18. Watson BO, Levenstein D, Greene JP, Gelinas JN, Buzsáki G. (2016); Multi-unit spiking activity recorded from rat frontal cortex (brain regions mPFC, OFC, ACC, and M2) during wake-sleep episode wherein at least 7 minutes of wake are followed by 20 minutes of sleep. Crcns.org. 10.6080/k02n506q.

19. F. Tatsuki, et al., Involvement of Ca(2+)-Dependent Hyperpolarization in Sleep Duration in Mammals. Neuron 90, 70–85 (2016).

20. K. Yoshida, et al., Leak potassium channels regulate sleep duration. Proc. Natl. Acad. Sci. U. S. A. 115, E9459–E9468 (2018).

21. L. B. Krone, et al., A role for the cortex in sleep-wake regulation. Nat. Neurosci. 24, 1210–1215 (2021).

22. G. H. Diering, et al., Homer1a drives homeostatic scaling-down of excitatory synapses during sleep. Science 355, 511–515 (2017).

23. Z. Wang, et al., Quantitative phosphoproteomic analysis of the molecular substrates of sleep need. Nature 558, 435–439 (2018).

24. F. Brüning, et al., Sleep-wake cycles drive daily dynamics of synaptic phosphorylation. Science 366 (2019).

25. A. A. Borbély, A two process model of sleep regulation. Hum. Neurobiol. 1, 195–204 (1982).

26. K. L. Ode, H. R. Ueda, Phosphorylation Hypothesis of Sleep. Front. Psychol. 11, 575328 (2020).

27. D. Tone, et al., Distinct phosphorylation states of mammalian CaMKIIβ control the induction and maintenance of sleep. PLoS Biol. 20, e3001813 (2022).

28. A. Hudmon, H. Schulman, Neuronal CA2+/calmodulin-dependent protein kinase II: the role of structure and autoregulation in cellular function. Annu. Rev. Biochem. 71, 473–510 (2002).

29. C. Cirelli, C. M. Gutierrez, G. Tononi, Extensive and divergent effects of sleep and wakefulness on brain gene expression. Neuron 41, 35–43 (2004).

30. K. B. Hengen, M. E. Lambo, S. D. Van Hooser, D. B. Katz, G. G. Turrigiano, Firing rate homeostasis in visual cortex of freely behaving rodents. Neuron 80, 335–342 (2013).

31. V. V. Vyazovskiy, et al., Cortical firing and sleep homeostasis. Neuron 63, 865–878 (2009).

32. V. V. Vyazovskiy, C. Cirelli, M. Pfister-Genskow, U. Faraguna, G. Tononi, Molecular and electrophysiological evidence for net synaptic potentiation in wake and depression in sleep. Nat. Neurosci. 11, 200–208 (2008).

33. L. de Vivo, et al., Ultrastructural evidence for synaptic scaling across the wake/sleep cycle. Science 355, 507–510 (2017).

34. S. J. Aton, et al., Mechanisms of sleep-dependent consolidation of cortical plasticity. Neuron 61, 454–466 (2009).

35. M. Massimini, F. Amzica, Extracellular calcium fluctuations and intracellular potentials in the cortex during the slow sleep oscillation. J. Neurophysiol. 85, 1346–1350 (2001).

36. R. Havekes, S. J. Aton, Impacts of Sleep Loss versus Waking Experience on Brain Plasticity: Parallel or Orthogonal? Trends Neurosci. 43, 385–393 (2020).

37. A. Suppermpool, D. G. Lyons, E. Broom, J. Rihel, Sleep pressure modulates single-neuron synapse number in zebrafish. Nature 629, 639–645 (2024).

38. R. M. Mulkey, S. Endo, S. Shenolikar, R. C. Malenka, Involvement of a calcineurin/inhibitor-1 phosphatase cascade in hippocampal long-term depression. Nature 369, 486–488 (1994).

39. J. Tomita, et al., Pan-Neuronal Knockdown of Calcineurin Reduces Sleep in the Fruit Fly, Drosophila melanogaster. J. Neurosci. 31, 13137–13146 (2011).

40. Y. Wang, et al., Post-synaptic competition between calcineurin and PKA regulates mammalian sleep-wake cycles. bioRxiv 2023.12.21.572751 (2023).

41. A. M. Strijkstra, S. Daan, Dissimilarity of slow-wave activity enhancement by torpor and sleep deprivation in a hibernator. Am. J. Physiol. 275, R1110–7 (1998).

42. C. G. von der Ohe, C. Darian-Smith, C. C. Garner, H. C. Heller, Ubiquitous and temperature-dependent neural plasticity in hibernators. J. Neurosci. 26, 10590–10598 (2006).

43. J.-G. He, H.-Y. Zhou, F. Wang, J.-G. Chen, Dysfunction of glutamatergic synaptic transmission in depression: Focus on AMPA receptor trafficking. Biol. Psychiatry Glob. Open Sci. 3, 187–196 (2023).

44. Y. Wei, G. P. Krishnan, M. Komarov, M. Bazhenov, Differential roles of sleep spindles and sleep slow oscillations in memory consolidation. PLoS Comput. Biol. 14, e1006322 (2018).

45. T. Tadros, M. Bazhenov, Role of Sleep in Formation of Relational Associative Memory. J. Neurosci. 42, 5330–5345 (2022).

46. J. F. A. Poulet, C. C. H. Petersen, Internal brain state regulates membrane potential synchrony in barrel cortex of behaving mice. Nature 454, 881–885 (2008).

47. I. Meir, Y. Katz, I. Lampl, Membrane Potential Correlates of Network Decorrelation and Improved SNR by Cholinergic Activation in the Somatosensory Cortex. J. Neurosci. 38, 10692–10708 (2018).

48. Z. Brzosko, S. B. Mierau, O. Paulsen, Neuromodulation of Spike-Timing-Dependent Plasticity: Past, Present, and Future. Neuron 103, 563–581 (2019).

49. L. Aitchison, et al., Synaptic plasticity as Bayesian inference. Nat. Neurosci. 24, 565–571 (2021).

50. D. Festa, A. Aschner, A. Davila, A. Kohn, R. Coen-Cagli, Neuronal variability reflects probabilistic inference tuned to natural image statistics. Nat. Commun. 12, 3635 (2021).

51. G. Shimizu, K. Yoshida, H. Kasai, T. Toyoizumi, Computational roles of intrinsic synaptic dynamics. Curr. Opin. Neurobiol. 70, 34–42 (2021).

52. G. Buzsáki, K. Mizuseki, The log-dynamic brain: how skewed distributions affect network operations. Nat. Rev. Neurosci. 15, 264–278 (2014).

53. M. Anisimova, et al., Spike-timing-dependent plasticity rewards synchrony rather than causality. Cereb. Cortex 33, 23–34 (2022).

54. W. M. Vanderheyden, J. R. Gerstner, A. Tanenhaus, J. C. Yin, P. J. Shaw, ERK phosphorylation regulates sleep and plasticity in Drosophila. PLoS One 8, e81554 (2013).

55. S. J. Kim, et al., Kinase signalling in excitatory neurons regulates sleep quantity and depth. Nature 612, 512–518 (2022).

56. B. L. Sabatini, T. G. Oertner, K. Svoboda, The life cycle of Ca(2+) ions in dendritic spines. Neuron 33, 439–452 (2002).

57. H. W. Kessels, R. Malinow, Synaptic AMPA receptor plasticity and behavior. Neuron 61, 340–350 (2009).

58. D. Levenstein, G. Buzsáki, J. Rinzel, NREM sleep in the rodent neocortex and hippocampus reflects excitable dynamics. Nat. Commun. 10, 2478 (2019).

59. P. Franken, D. Chollet, M. Tafti, The homeostatic regulation of sleep need is under genetic control. J. Neurosci. 21, 2610–2621 (2001).

