## Supplementary Materials for "Boundary conditions for synaptic homeodynamics during the sleep-wake cycle"

**This PDF file includes:**

Supporting information text
Figures. S1 to S20
Tables S1 to S8
References

### Supporting Information Text

#### Parameter search for synaptic learning rules

We selected parameter values from uniform distributions within the ranges of values (1) (**Table S2**). Parameter sets for Hebbian and STDP were collected in the conditions where $\theta_p > \theta_d$ because most of them did not appear in $\theta_p < \theta_d$. In the same way, parameter sets for Anti-Hebbian and Anti-STDP were collected in $\theta_p < \theta_d$. The synaptic changes in different time lags were calculated analytically according to the way presented by previous reports (1). Neurons were stimulated at 1 Hz for 60 seconds. First, scaling parameters $\beta_{NMDA}$ $and$ $\beta_{VGCC}$ in the equations (*Eq.* 3 and 8, *Materials and Methods* in the main text) were calculated so that the mean amplitudes of $C_{pre}$ and $C_{post}$ ($MC_{pre}$ and $MC_{post}$) during 2-3 seconds satisfy $MC_{pre}$ = 0.7 μM and $MC_{post}$ =1.4 μM because this kind of relationship holds in the experimental studies (2). We considered that synaptic weights did not change if spike-pairs had large time differences. Specifically, the ratio of $\gamma_p$ and $\gamma_d$ was calculated and the value of $\gamma_d$ was updated so that the potentiation and depression rate become equal when the lag time was equal to 100 milliseconds. The potentiation and depression rate were expressed as $\gamma_p\alpha_p$ and $\gamma_d\alpha_d$ respectively ( $\alpha_x = \frac{1}{T}\int_0^T \Theta[c(t) - \theta_x]\, dt,$ $T$ is the duration of the stimulation protocol) (1). Then, the scaling parameters were calculated again in the parameter sets where $\gamma_d$was updated and synaptic changes were analytically calculated in each lag time (from -160 milliseconds to 160 milliseconds). To speed up the calculations, potentiation and depression rate were calculated during the 2-3 seconds and multiplied by 60. Gaussian curves that represent four different types of synaptic learning rules were described by the following equations with amplitude ($a_{lr}$, $b_{lr}$, $|a_{lr}| = |b_{lr}|$) and time constant ($\tau_{lr}$).

$$a_{lr} exp\left(-\left(\frac{x}{\tau_{lr}}\right)^2\right)\ (x \geq 0)$$

$$b_{lr} exp\left(-\left(\frac{x}{\tau_{lr}}\right)^2\right)\ (x < 0)$$

For each parameter set, the sum of squared errors (SSE) between analytical solutions and fitting gaussian curves were calculated and the parameter set was collected if the SSE was less than the threshold (The threshold was 0.45 in $a_{lr} = 0.9$, $\tau_{lr} = 50$ and 0.25 in all other cases of **Fig 1*E*** and **Fig. S1 *A-E***. The threshold was 0.6 when searching for synaptic learning rules in Hodgkin–Huxley-based network models).

**Noise term and comparison of simulation and analytical results**

The noise term in the equation (*Eq.*1, *Materials and Methods* in the main text) was expressed by the following equation.

$$Noise\ (t) = \sigma \sqrt{\frac{\pi\left[\Theta(c(t)-\theta_d)+\Theta\left(c(t)-\theta_p\right)\right]z}{dt}}\ \eta(t)$$

When solving the differential equations, the noise term was calculated by above equation and added to the differential equation. *z* is the coefficient of noise, *dt* is a step size, and $\eta(t)$ is a Gaussian white noise process with unit variance density. We simulated the synaptic efficacy by the equation (*Eq.*1) for 1000 times and calculated the synaptic changes in different lag times according to the previous article (1). An analytical solution was compared to simulation results in different values of *z* in the simple model (**Fig. S17**) and *z* = 3.5 was used in the simulations with synaptic learning rules in this study.

**Generation of sleep and wake-like spike patterns**

We assumed that synchronization of firing was observed in NREM sleep states and desynchronization of firing was observed during wakefulness. Timestamps of wake-like spikes were obtained from ISI per spike, and those of sleep-like spikes were obtained from ISI per spike and duration of Up and Down states per burst. ISI per spike and durations of Up and Down states per burst were sampled from lognormal distributions with specific means and SDs. Spike patterns with a certain mean firing rate in **Figs. 1*H***, and **2*A-D*,** and **Figs. S3, S16*A***, and **S19** were generated by adjusting the mean Down-state duration. Spike patterns in **Figs. 2*E*** and **5*A***, and **Fig. S2** were generated by adjusting the mean ISI.

The relationships between mean and SD for the lognormal distributions were obtained based on previous in vivo recordings (3, 4) (**Fig. S18 *C-G***).

**Definition of lognormal distributions based on in vivo recordings**

The duration of Up states and Down states were sampled from the lognormal distributions because previous studies found that duration of Up states and Down states have lognormal distributions (3, 5). We assumed that ISI also has lognormal distributions on the same datasets (3, 4) (**Fig. S18 *A*** and ***B***). The spike data recorded from excitatory neurons (verified by cross-correlogram) of rats which had natural sleep-wake cycle from the previous studies (3, 4) were used in this study. To reduce the variables, liner regression analysis was performed on the ordinary logarithm of mean and SD of the duration of Up and Down states (**Fig. S18 *C*** and ***D***) and ISI (**Fig. S18 *E*** and ***F***), and the ordinary logarithm of the mean Up-state duration and mean Down-state durations (**Fig. S18*G***). Then, the SD of Up-state duration, Down-state duration, ISI in the Up states of sleep-like firing patterns and ISI in wake-like firing patterns were calculated by the equations: 0.35 × log10(*mean Up-state duration*) - 0.7, 0.25 × log10(*mean Down-state duration*) - 0.35, - 0.2 × log10(*mean ISI in the Up states of sleep*) + 0.95, and 0.03 × log10(*mean ISI in the state of wake*) + 0.65, respectively. The mean Down-state duration was calculated by the equation -0.7 × log10(*mean Up-state duration*) + 4.0.

**Conversion from spike patterns to voltage waveforms**

The timestamp data for spikes during sleep-like and wake -like firing patterns were converted to voltage waveforms. We defined several parameters when constructing voltage waveforms so that the timestamps of the spikes corresponded to the peaks of the membrane potential by referring to the experimental studies (6, 7) (**Table S3**). Voltage waveforms were generated by linear interpolation of the points for the membrane potential of peaks, after hyperpolarization and Up states. The difference of membrane potentials between Up states and Down states was 15 mV when calculating synaptic efficacy in **Figs. 1*H*, 2 *A-E*** and **5*A*,** and **Figs. S2, S3** and **S16**. The results in other values for the difference of membrane potentials are shown in **Fig. S19**.

**Hodgkin-Huxley-based network model**

A Hodgkin-Huxley-based network model was constructed based on the averaged neuron model in a previous study with some modifications (8). The equations of the network model are as follows.

$$CA\frac{dV}{dt} = -A\left(I_L(V_{post}) + I_{Na}(V_{post}, h_{na}) + I_K(V_{post}, n_k) + I_A(V_{post}, h_A) + I_{KS}(V_{post}, m_{KS}) + I_{Ca}(V_{post})\right.$$

$$\left. + I_{KCa}(V_{post}, [Ca^{2+}]) + I_{NaP}(V_{post}) + I_{AR}(V_{post})\right) - I_{NMDA}(V_{post}, s_{NMDA}, x_{NMDA})$$

$$- I_{AMPA}(V_{post}, s_{AMPA}) - I_{GABA}(V_{post}, s_{GABA})$$

$$\frac{dh_{na}}{dt} = 4(\alpha_h(V_{post})(1 - h_{Na}) - \beta_h(V_{post})h_{Na})$$

$$\frac{dn_k}{dt} = 4(\alpha_n(V_{post})(1 - n_k) - \beta_n(V_{post})n_k)$$

$$\frac{dh_A}{dt} = (h_{A\infty}(V_{post}) - h_A)/\tau_{hA}$$

$$\frac{dm_{KS}}{dt} = (m_{KS\infty}(V_{post}) - m_{KS})/\tau_{mKS}(V_{post})$$

$$\frac{ds_{AMPA}}{dt} = a_{AMPA}f(V_{pre}) - \frac{s_{AMPA}}{\tau_{AMPA}}$$

$$\frac{ds_{NMDA}}{dt} = a_{sNMDA}(1 - s_{NMDA}) - \frac{s_{NMDA}}{\tau_{sNMDA}}$$

$$\frac{dx_{NMDA}}{dt} = a_{xNMDA}f(V_{pre}) - \frac{x_{NMDA}}{\tau_{xNMDA}}$$

$$\frac{ds_{GABA}}{dt} = a_{GABA}f(V_{pre}) - \frac{s_{GABA}}{\tau_{GABA}}$$

$$\frac{d[Ca^{2+}]}{dt} = -\alpha_{Ca}\left(AI_{Ca}(V_{post})\right) - \frac{[Ca^{2+}]}{\tau_{Ca}}$$

$$\frac{d[Ca_{syn}^{2+}]}{dt} = -\alpha_{Ca}(AI_{Ca}(V_{post})\beta_{VGCC} + I_{NMDA}(V_{post}, s_{NMDA}, x_{NMDA})\beta_{NMDA}) - \frac{[Ca_{syn}^{2+}]}{\tau_{Ca-syn}}$$

, where $C$ is the membrane capacitance, $A$ is the area of a single neuron, $V$ is the membrane potential, $[Ca^{2+}]$ is the intracellular calcium concentration, and $IX$ ($X$: each ion channel) denotes the current of each ion channel. Each function is listed below.

$$I_L(V_{post}) = g_L(V_{post} - V_L)$$

$$I_{Na}(V_{post}, h_{na}) = g_{Na} m_{Na\infty}^3 (V_{post}) h_{Na} (V_{post} - V_{Na})$$

$$m_{Na\infty}(V_{post}) = \alpha_m(V_{post})/(\alpha_m(V_{post}) + \beta_m(V_{post}))$$

$$\alpha_m(V_{post}) = 0.1(V_{post} + 33)/[1 - exp(-(V_{post} + 33)/10)]$$

$$\beta_m(V_{post}) = 4exp(-(V_{post} + 53.7)/12)$$

$$\alpha_h(V_{post}) = 0.07exp(-V_{post} + 50)/10)$$

$$\beta_h(V_{post}) = 1/[1 + exp(-(V_{post} + 20)/10)]$$

$$I_K(V_{post}, n_k) = g_K n_K^4 (V_{post} - V_K)$$

$$\alpha_n(V_{post}) = 0.01(V_{post} + 34)/[1 - exp(-(V_{post} + 34)/10)]$$

$$\beta_n(V_{post}) = 0.125exp(-(V_{post} + 44)/25)$$

$$I_A(V_{post}, h_A) = g_A m_{A\infty}^3 (V_{post}) h_A (V_{post} - V_K)$$

$$m_{A\infty}(V_{post}) = 1/[1 + exp(-(V_{post} + 50)/2)]$$

$$h_{A\infty}(V_{post}) = 1/[1 + exp((V_{post} + 80)/6)]$$

$$I_{KS}(V_{post}, m_{KS}) = g_{KS} m_{KS} (V_{post} - V_K)$$

$$m_{KS\infty}(V_{post}) = 1/[1 + exp(-(V_{post} + 34)/6.5)]$$

$$\tau_{mKS}(V_{post}) = 8/[exp(-(V_{post} + 55)/30) + exp((V_{post} + 55)/30)]$$

$$I_{Ca}(V_{post}) = g_{Ca} m_{Ca\infty}^2 (V)(V_{post} - V_{Ca})$$

$$m_{Ca\infty}(V_{post}) = 1/[1 + exp(-(V_{post} + 20)/9)]$$

$$I_{KCa}(V_{post}, [Ca^{2+}]) = g_{KCa} m_{KCa\infty}([Ca^{2+}])(V_{post} - V_K)$$

$$m_{KCa\infty}([Ca^{2+}]) = 1/[1 + (K_D/[Ca^{2+}])^{3.5}]$$

$$I_{NaP}(V_{post}) = g_{NaP} m_{NaP\infty}^3 (V_{post})(V_{post} - V_{Na})$$

$$m_{NaP\infty}(V_{post}) = 1; [1 + exp(-(V_{post} + 55.7)/7.7)]$$

$$I_{AR}(V_{post}) = g_{AR} h_{AR\infty}(V_{post})(V_{post} - V_K)$$

$$h_{AR\infty}(V_{post}) = 1/[1 + exp((V_{post} + 75)/4)]$$

$$f(V_{post}) = 1/[1 + exp(-(V_{post} - 20)/2)]$$

$$I_{AMPA}(V_{post}, s_{AMPA}) = g_{AMPA}s_{AMPA}(V_{post} - V_{AMPA})$$

$$I_{NMDA}(V_{post}, s_{NMDA}, x_{NMDA}) = g_{NMDA}s_{NMDA}(V_{post} - V_{NMDA})$$

$$I_{GABA}(V_{post}, s_{GABA}) = g_{GABA}s_{GABA}(V_{post} - V_{GABA})$$

The constant values are listed in **Table S4**. Intrinsic (non-synaptic) ion currents (e.g. $I_{Na}$) should be multiplied by 10 to adjust its unit to nanoampere (nA) when the numerical values listed in **Table S5** are directly used in the numerical simulations.

The network model has two $Ca^{2+}$ compartments of cell body and synapse following the facts that the time constant of $Ca^{2+}$ in synapses is very short (about 15 milliseconds) (2) while a larger $Ca^{2+}$ time constant (about 50-1000 milliseconds) is necessary to induce SWO (8). We did not consider the influx of $Ca^{2+}$ from spines to a cell body taking into account the immediate uptake of $Ca^{2+}$ by intracellular buffers (9). The number of excitatory and inhibitory synapses per neuron were sampled from lognormal distributions, respectively. Connections between inhibitory neurons were not considered because we did not include this type of connection in parameter search and bifurcation analysis in a single neuron. The mean = 2 and SD = 0.01 was adopted for lognormal distributions of the number of excitatory and inhibitory synapses per neuron in **Figs. 3** and **4**, in which excitatory neurons had both two excitatory and inhibitory synapses and inhibitory neurons had only two excitatory synapses. The conductance of AMPAR and NMDAR or the conductance of GABAR were divided by the average number of excitatory or inhibitory synapses per neuron respectively in the simulations of Hodgkin-Huxley-based network models.

**Parameter search for SWO and bifurcation analysis in Hodgkin-Huxley-based network models**

Parameter search for SWO and bifurcation analysis for a single neuron were based on the previous article (8). First, we randomly generated parameter sets from a large parameter space and searched for parameter sets that yield firing patterns of SWO. The ranges of parameters were defined to include biophysically reasonable values. The conductance of intrinsic (non-synaptic) channels in the soma and axon ($g_L$, $g_{Na}$, $g_K$, $g_A$, $g_{KS}$, $g_{Ca}$, $g_{KCa}$, $g_{NaP}$, $g_{AR}$) and extrinsic (synaptic) channels in the dendrite ($g_{AMPA}$, $g_{NMDA}$, $g_{GABA}$) were generated by selecting parameters from an exponential distribution bounded to the interval $10^{-2}$-$10^{2}$ mS/cm$^2$ and $10^{-3}$-$10^{1}$ μS, respectively. A random parameter search for time constant of intracellular $Ca^{2+}$ ($\tau_{Ca}$) were also conducted in the interval $10^1$-$10^3$ milliseconds. The parameter sets that yielded a steady-state and periodic solution with real values were evaluated. The major frequency of the oscillatory behavior was analyzed by fast Fourier transform. Additionally, we assessed the detailed structure of the oscillations by counting the number of spikes; the number of spikes was determined as half the number of times the membrane potential crossed -20 mV. Solutions in which the maximum of membrane potential were more than 200 mV or the minimum of membrane potential were less than -200 mV were eliminated at this point. Based on these characteristics, the solutions were classified into six categories: “Resting” (spike numbers per second < 0.6, peak frequency <= 0.5 Hz, maximum membrane potential < -30 mV or minimum membrane potential > -30 mV), “SWO” (peak frequency >=0.5 Hz and 2 × peak frequency <= spike number per second < 30), “SWO with high frequency” (peak frequency >=0.5 Hz, spike number per second >= 2 × peak frequency and spike number per second >= 30), “AWAKE” (peak frequency >=0.5 Hz, spike number per second <= 2 × peak frequency and spike number per second < 30), “AWAKE with high frequency” (peak frequency >=0.5 Hz, 30 <spike number per second <= 2 × peak frequency and spike number per second < 100) and “EXCLUDED” (others not classified above).

Second, we conducted bifurcation analysis in parameter sets that yield SWO or SWO with high frequency. In the bifurcation analysis, channel or receptor conductance was gradually changed from $10^{-2}$ to $10^{2}$ times its original value by 100 steps (for the bifurcation by the pre-synaptic mechanism, $a_{AMPA}$, $a_{xNMDA}$, $a_{GABA}$ were gradually changed from $10^{-2}$ to $10^{1}$ times its original value by 100 steps), and the solutions were automatically classified into six categories according to the criteria described above. We selected parameter sets that including the bifurcation from “AWAKE” to “SWO” and satisfying minimum membrane potential of “SWO” + 5 mV < minimum membrane potential of “AWAKE” in continuously varying conductance. In the parameter search and bifurcation analysis for a single neuron, the simulations were conducted for 6 seconds, and the results of 1-6 seconds were analyzed. Integration was performed with the following initial values in each simulation of parameter search and bifurcation analysis for a single neuron: $V$ = -45 mV, $h_{Na}$ = 0.045 (unitless), $n_K$ = 0.54 (unitless), $h_A$ = 0.045 (unitless), $m_{KS}$ = 0.34 (unitless), $[Ca^{2+}]$ = 1 μM, $s_{AMPA}$ = 0.01 (unitless), $s_{NMDA}$ = 0.01 (unitless), $x_{NMDA}$ = 0.01 (unitless), $s_{GABA}$ = 0.01 (unitless).

Then, the network model of 80 neurons was constructed with E:I ratio of 4:1 and bifurcation analysis for network models was conducted. In the first bifurcation analysis for network models, channel or receptor conductance was gradually changed from $10^{-2}$ to $10^{2}$ times its original value by 17 steps (for the bifurcation by the pre-synaptic mechanism, $a_{AMPA}$, $a_{xNMDA}$, $a_{GABA}$ were gradually changed from $10^{-2}$ to $10^{1}$ times its original value by 16 steps) and the simulations were conducted for 2.5 seconds. The membrane potential of all neurons in a network model were analyzed and classified into three categories based on the above criteria: “Sleep” (classified into “SWO” or “SWO with high frequency”), “Wake” (classified into “AWAKE” or “AWAKE with high frequency”) and “Others” (classified into other categories).

For the evaluation of the degree of synchronization and desynchronization, coefficient of variance (CV) for total spike counts per 50 milliseconds (sleep score) was calculated. More than 1000 parameter sets that bifurcated from wake-like patterns as a network (defined by sleep scores < 1.0 and the percentage of “Wake” neurons > 30 %) to wake-like patterns as a network (defined by sleep score >= 1.3 and the percentage of “Sleep” neurons > 30 %) were selected in each bifurcation model. Initial values in a network model were randomly assigned for $V$, $h$, $h_{Na}$, $n_K$, $h_A$ and $m_{KS}$ and other variables were the same as those of bifurcation analysis in a single neuron.

We conducted a second bifurcation analysis for the parameter sets collected by the first bifurcation analysis. In the second bifurcation analysis, the simulations were conducted for 5 seconds and the conductance was gradually changed from $10^{-2}$ to $10^{1.5}$ times its original value by 36 steps (for the bifurcation by the pre-synaptic mechanism, $a_{AMPA}$, $a_{xNMDA}$, $a_{GABA}$ were gradually changed from $10^{-2}$ to $10^{0.8}$ times its original value by 29 steps). The results were analyzed in the same way as the first bifurcation analysis and manually checked. The parameter sets including a stable sleep-like and wake-like firing patterns as a network with those mean firing rates that were close to each other (the difference is less than 2.0 Hz, called a representative sleep-like and wake-like state respectively) were selected.

We also constructed network models with different connection patterns because real connections of cortical networks are non-random lognormal-like connections (10). The network models had different SDs and means of the lognormal distributions for the number of synapses per neuron from the original model (**Fig. S10 *A*** and ***D***). Parameter sets that showed transitions from wake-like to sleep-like firing patterns after the first bifurcation analysis were selected. In the second bifurcation analysis, the ranges of the conductance or coefficient were from $10^{(\boldsymbol{wv}-0.7)}$ to $10^{(\boldsymbol{sv}+0.7)}$ times its original value by (*sv* – *wv* + 1.4) × 10 steps (*wv* and *sv* were the values for the representative sleep-like and wake-like states as a network in the original model, respectively).

**Parameter search for synaptic learning rules in Hodgkin-Huxley-based network models**

After the second bifurcation analysis, we analytically calculated the synaptic changes and searched for parameter sets representing specific learning rules in the same way as simple models. When using learning rules in network models, the equation (*Eq.* 4, *Materials and Methods* in main text) was replaced by the following equation.

$$I_{Ca\_NMDA} = g_{NMDA} s_{NMDA} (V_{post} - V_{Ca})$$

In details, we used a 4-second time series data for membrane potentials of a single neuron in wake-like states and duplicate the data with a short delay in each time lag (**Fig. S6**). Parameter sets for synaptic learning rules were randomly generated and the periods that post-synaptic $Ca^{2+}$ exceeded $\theta_p$ or $\theta_d$in 3 seconds (from one to four seconds) were calculated respectively. Then, the periods spent above each threshold were multiplied by 20 to obtain the total values for 60 seconds for the purpose of speeding up the calculations, considering that the results would not change so much if time series data for membrane potentials were stable. Therefore, we selected time series data for membrane potentials of a representative neuron that had the minimum CV for firing rates in 1-second non-overlapping windows. Synaptic changes in each time lag were analytically calculated and the parameter sets of which the SSE between analytical results and the Gaussian fitting curves was less than 0.6 were selected. The ranges of $\theta_p$ and $\theta_d$ were limited to [0.8,1.6] and [0.5, 1.0] when searching STDP and limited to [0.5, 1.0] and [0.8,1.6] when searching Anti-STDP, respectively. The ranges of other parameters are shown in **Table S2**. The same parameter sets for synaptic learning rules were applied to all excitatory synapses in a network model.

**Obtaining values for scaling parameters and normalization of thresholds for synaptic learning rules**

Values for scaling parameters ($\beta_{NMDA}$ and $\beta_{VGCC}$) were calculated when we compare synaptic efficacy between sleep-like and wake-like firing patterns in Hodgkin-Huxley-based network models according to the following procedures. $\beta_{NMDA}$ was set to be 1.0 and $\beta_{VGCC}$ was obtained by satisfying $2MC_{pre} = MC_{post}$ based on the results of preliminary simulations as conducted in simple model. $MC_{pre}$ and $MC_{post}$ were the mean amplitudes of $C_{pre}$ and $C_{post}$ in a 10-seconds' simulation of wake-like patterns respectively, which were implemented in $\beta_{NMDA} = \beta_{VGCC} = 1.0$ and without learning rules (In the model with sleep-wake dynamics, $MC_{pre}$ and $MC_{post}$ were the mean amplitudes of $C_{pre}$ and $C_{post}$ during wake-like periods in simulations for 15 seconds in **Figs. S11, S13** and **S14** or 25 seconds in **Fig. 4** and **Fig.**

**S12**). $\theta_p$ and $\theta_d$ of synaptic learning rules were normalized by $MC_{pre}$ during wake-like patterns as the following equation.

$$\theta_x' = \frac{\theta_x \times MC_{pre}}{0.7} \,(x = p \; or \; d)$$

**Evaluation of synchronization and desynchronization in Hodgkin-Huxley-based network models**

For the evaluation of the degree of synchronization and desynchronization, CV for total spike counts per 50-milliseconds' window without overlaps was calculated (Sleep score). Sleep scores were obtained from the waveforms of membrane potentials in all neurons for 2.5 seconds or 5.0 seconds in the first and second bifurcation analysis, respectively (**Fig. 3** ***B-F***). In simulations with sleep-wake dynamics (**Fig. 4** ***C-F***), sleep scores were calculated continuously in 5-seconds' window with 500-milliseconds' overlaps. For the statistical tests, 1,000,000 sets of desynchronized spike trains for 2.5 seconds or 5 seconds (80 neurons, step size is 0.01) with the frequency of 0.5-15 Hz were randomly generated as the same way as in **Fig. 1** ***G*** and ***H*** and sleep scores were calculated in each dataset. A sleep score of the data was compared with this distribution and regarded as sleep-like states if it was more than the value with $p < 0.01$. The distribution of sleep scores in generated desynchronized spike trains and sleep scores with $p < 0.01$ in different conditions are shown in **Fig. S20**.

**Calculation of synaptic efficacy under synaptic learning rules in Hodgkin–Huxley-based network models including stimulation during the wakefulness**

The construction of the network model was the same as **Figs. 3** and **4**. In **Fig. S15**, excitatory neurons were divided into three groups by their indices: group **1** and **2** (20 neurons in each group) are the groups for stimulated neurons and group **3** (24 neurons) is for unstimulated neurons. First, all neurons fire with desynchronized wake-like firing patterns. Then, neurons in group **1** and **2** were stimulated simultaneously at the end of wake-like states. The stimulation was optimized by changing its waveforms and rates so that the potentiation of synaptic efficacy between stimulated groups are observed under STDP during the stimulation protocol. After the stimulation, all neurons fire with synchronized sleep-like firing patterns. Mean of synaptic efficacy every time step and ratio of mean synaptic efficacy after to before sleep-like firing patterns were calculated and compared between stimulated neurons and unstimulated neurons.

**Software and Numerical calculations**

We used Python (version 3.8) with these following libraries: jupyter, matplotlib, numpy, pandas, scikit-learn, statannot, numba, seaborn, scipy, and vistats.

All simulations were conducted using GPGPU (NVIDIA RTX 3090, A5000 or A6000), C++17 and CUDA (version 12.0). except the simulations of parameter search for synaptic learning rules or parameter search for SWO and bifurcation analysis in a single neuron. Calculations were implemented by using a forth-order Runge-Kutta method. Time steps were 0.1 milliseconds in parameter search for synaptic learning rules, 0.05 milliseconds in calculations of synaptic efficacy in simple model, and 0.00005-0.05 milliseconds in calculations of synaptic efficacy in Hodgkin–Huxley-based network models, respectively.

#### Supplementary Figures

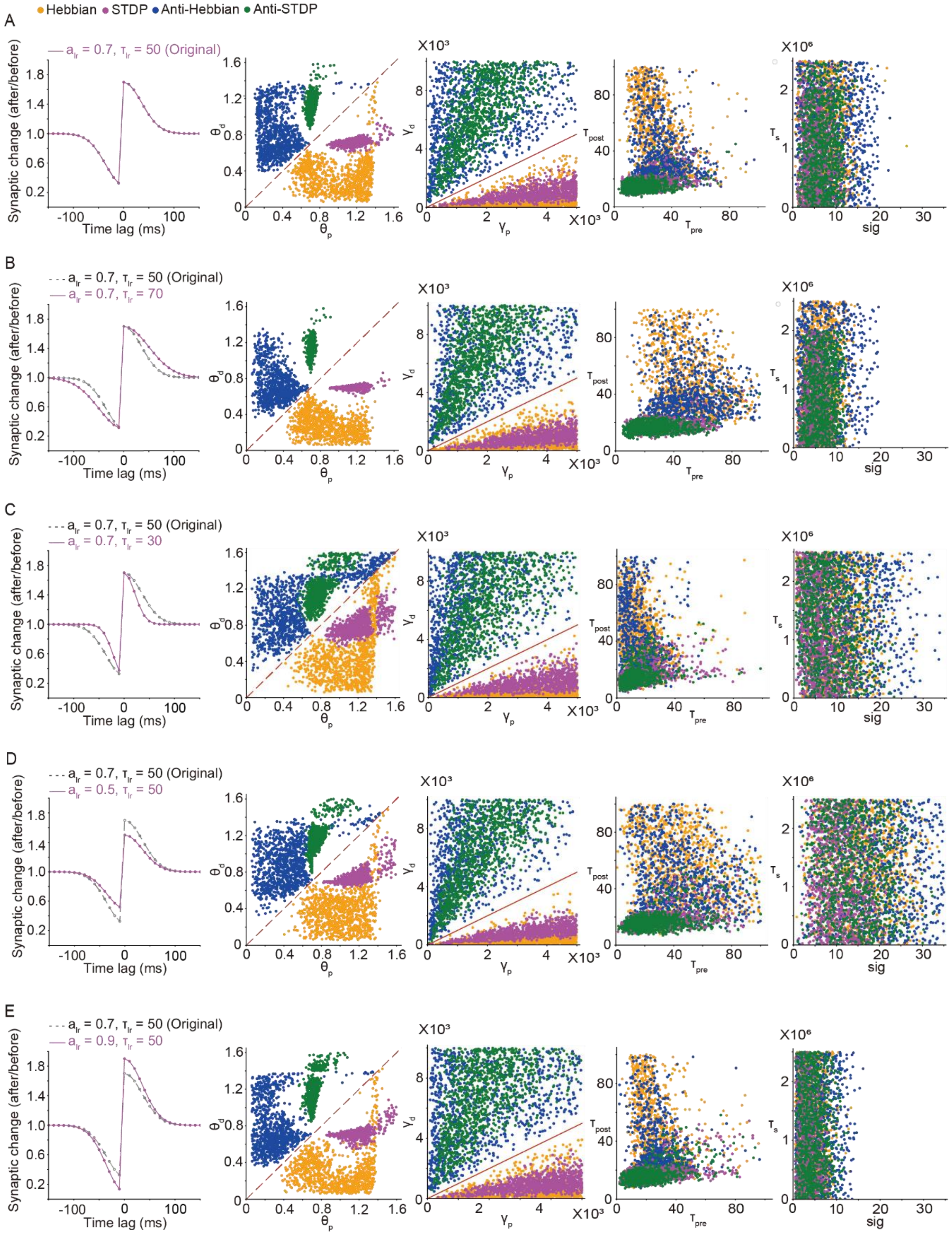

**Fig. S1. Distributions of parameter sets for synaptic learning rules in different fitting curves, related to Fig. 1**

Distributions for parameter sets for four types of synaptic learning rules obtained by different fitting curves.

(A) The original one. The plots in axes of thresholds and amplitudes were the same as in **Fig. 1*F***.

(B-E) Parameter sets were collected by fitting to gaussian curves with different amplitudes ($a_{lr}$) and time constants ($\tau_{lr}$) from original one. The ranges of parameters are shown in **Table S2**.

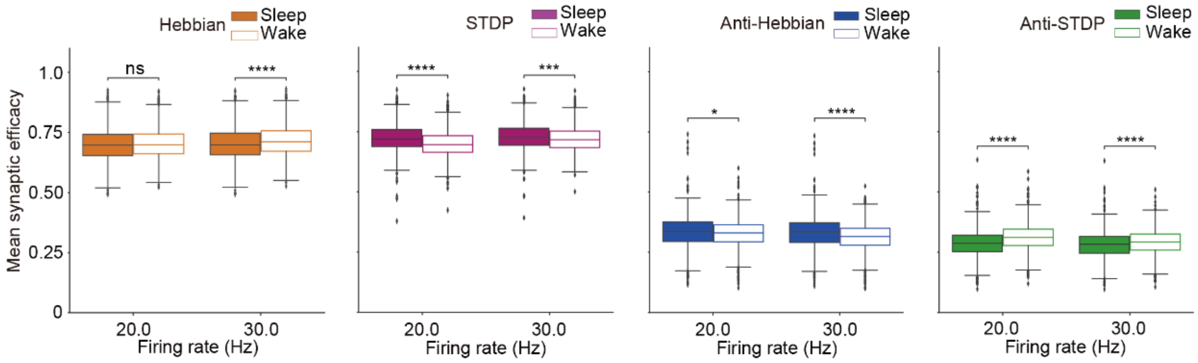


**Fig. S2. Mean synaptic efficacy at higher mean firing rates, related to Fig. 1**

Box plots for mean synaptic efficacy under four different types of synaptic learning rules at higher mean firing rates ($n$ = 1000 for each firing rate, $n$ represents the number of synaptic learning rules). The parameter sets for synaptic learning rules were the same as in **Fig. 1*H***. The sleep-like firing patterns were generated by sampling from the lognormal distributions for Up-state duration and Down-state duration (log10(*mean Up-state duration*) = 2.7 and log10(*mean Down-state duration*) = 2.7, SD was calculated according to the linear regression analysis based on in vivo data (**Fig. S18**)). The whiskers above and below of box plots show minimal to maximal values. The box extends from the 25th to the 75th percentile and the middle line indicates the median. * p < 0.05, ** p < 0.01, *** p < 0.001, **** p < 0.0001, Welch's t-test was applied.

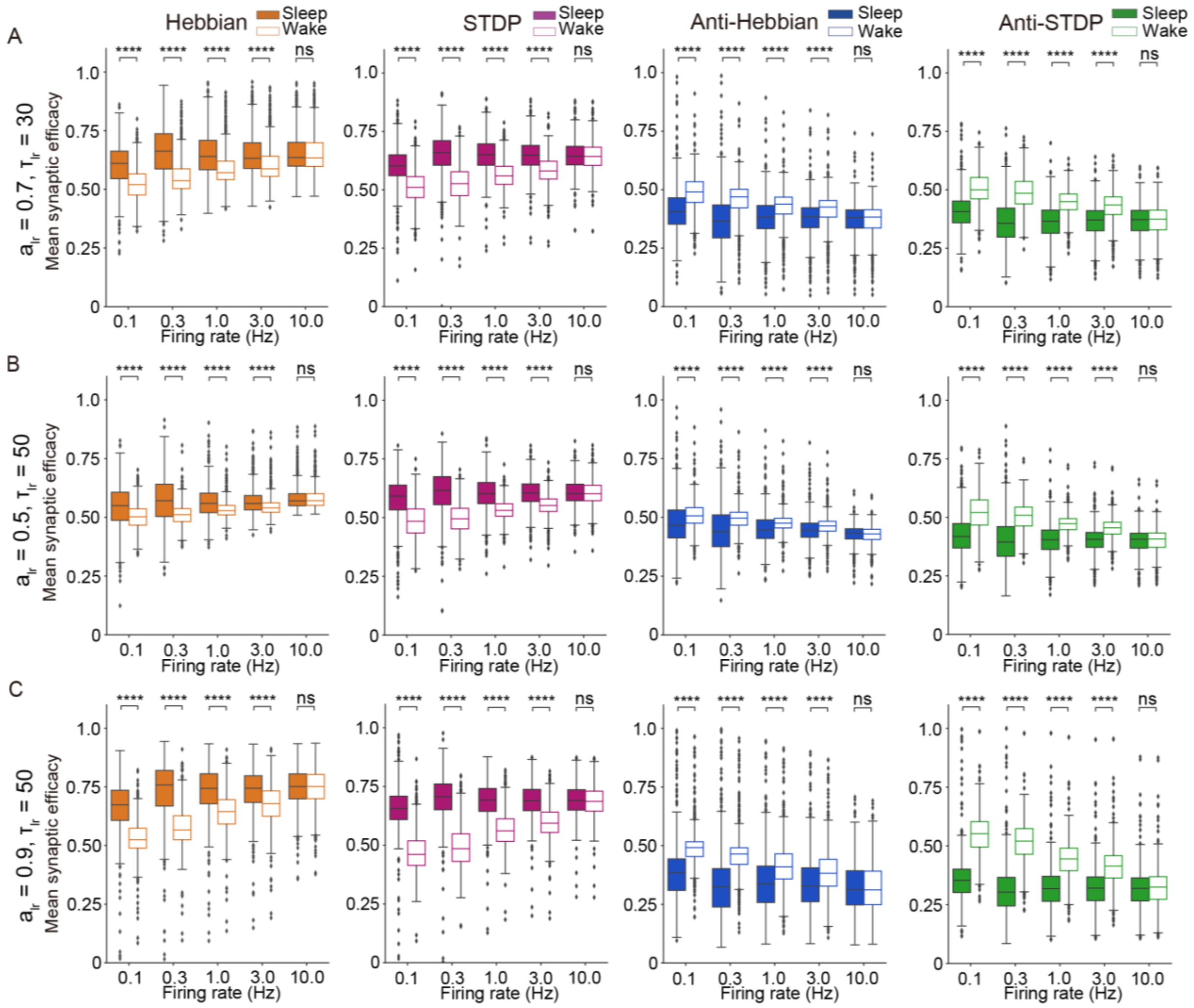


**Fig. S3. Mean synaptic efficacy in different fitting curves, related to Fig. 2**

(A-C) Box plots for mean synaptic efficacy under synaptic learning rules in different parameters for fitting curves (*n* = 1000 for each firing rate, *n* represents the number of synaptic learning rules). The results of $a_{lr} = 0.7, \tau_{lr} = 30$ (A), $a_{lr} = 0.5, \tau_{lr} = 50$ (B), and $a_{lr} = 0.9, \tau_{lr} = 50$ (C) are shown. The parameter sets for synaptic learning rules were the same as in **Fig. S1**. The sleep-like firing patterns were the same as in **Fig. 1*H***. The whiskers above and below of box plots show minimal to maximal values. The box extends from the 25th to the 75th percentile and the middle line indicates the median. * $p < 0.05$, ** $p < 0.01$, *** $p < 0.001$, **** $p < 0.0001$, Welch's t-test was applied.

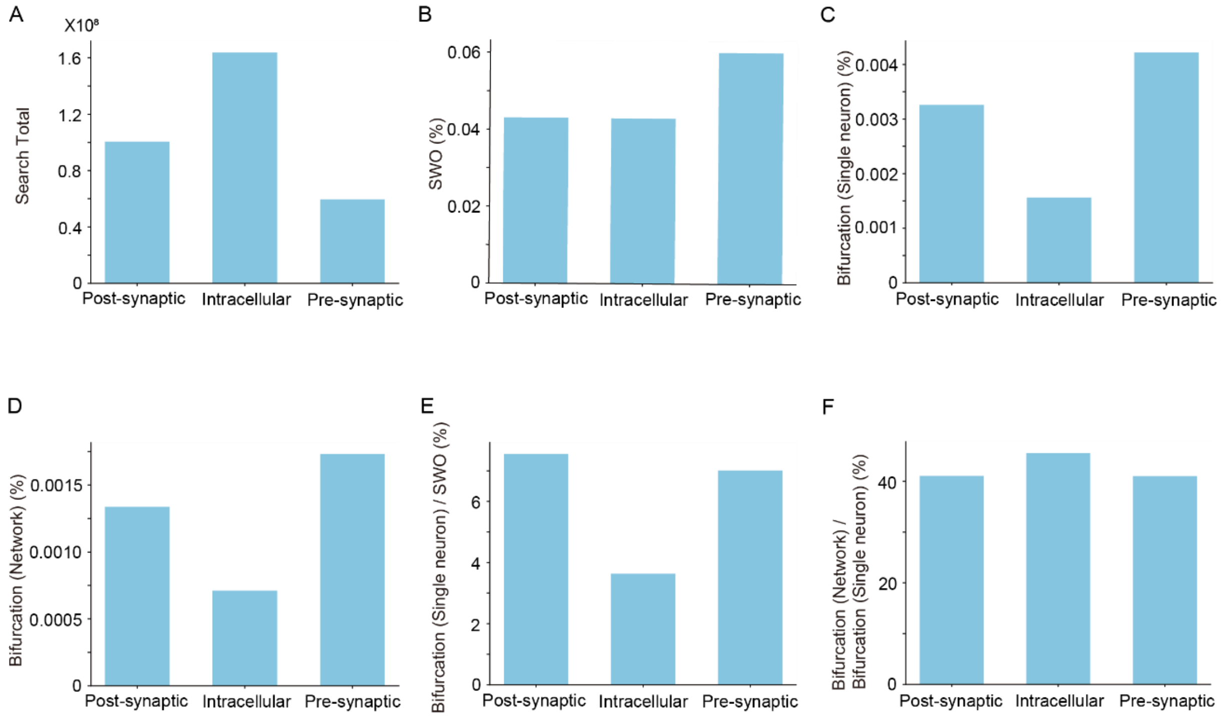


**Fig. S4. Results of parameter search for SWO and bifurcation analysis, related to Fig. 3**

(A-F) The results of parameter search for SWO and bifurcation analysis in single neurons and network models are shown by three types of bifurcation models.

The longitudinal axes represent the number of total parameter sets searched (A), the percentage of parameter sets generating SWO (B), the percentage of parameter sets bifurcating from wake-like to sleep-like firing patterns in a single neuron (bifurcation (single neuron)) (C), the percentage of parameter sets bifurcating from wake-like to sleep-like firing patterns as a network (bifurcation (network)) (D), ratio of bifurcation (single neuron) to SWO (E), ratio of bifurcation (network) to bifurcation (single neuron) (F).

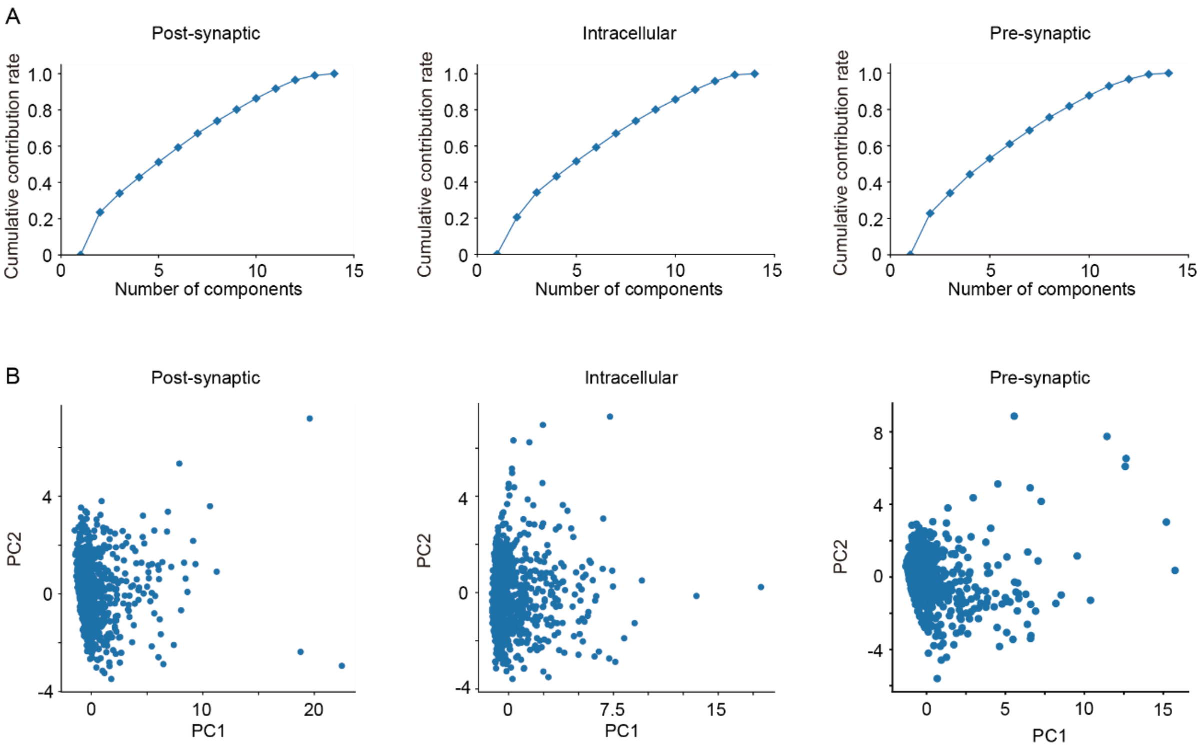


**Fig. S5. Clustering of the results of bifurcation analysis, related to Fig. 3**

Principal component analysis (PCA) was conducted for the parameter sets that bifurcated form sleep-like to wake-like firing patterns as a network obtained in bifurcation analysis in each model.

(A) Cumulative contribution ratio of eigen values when PCA applied to the parameter sets.

(B) Projection of the parameter sets onto their first two principal components in each model (n = 1344, 1202, and 1092 in the post-synaptic, intracellular and pre-synaptic bifurcation models, respectively).

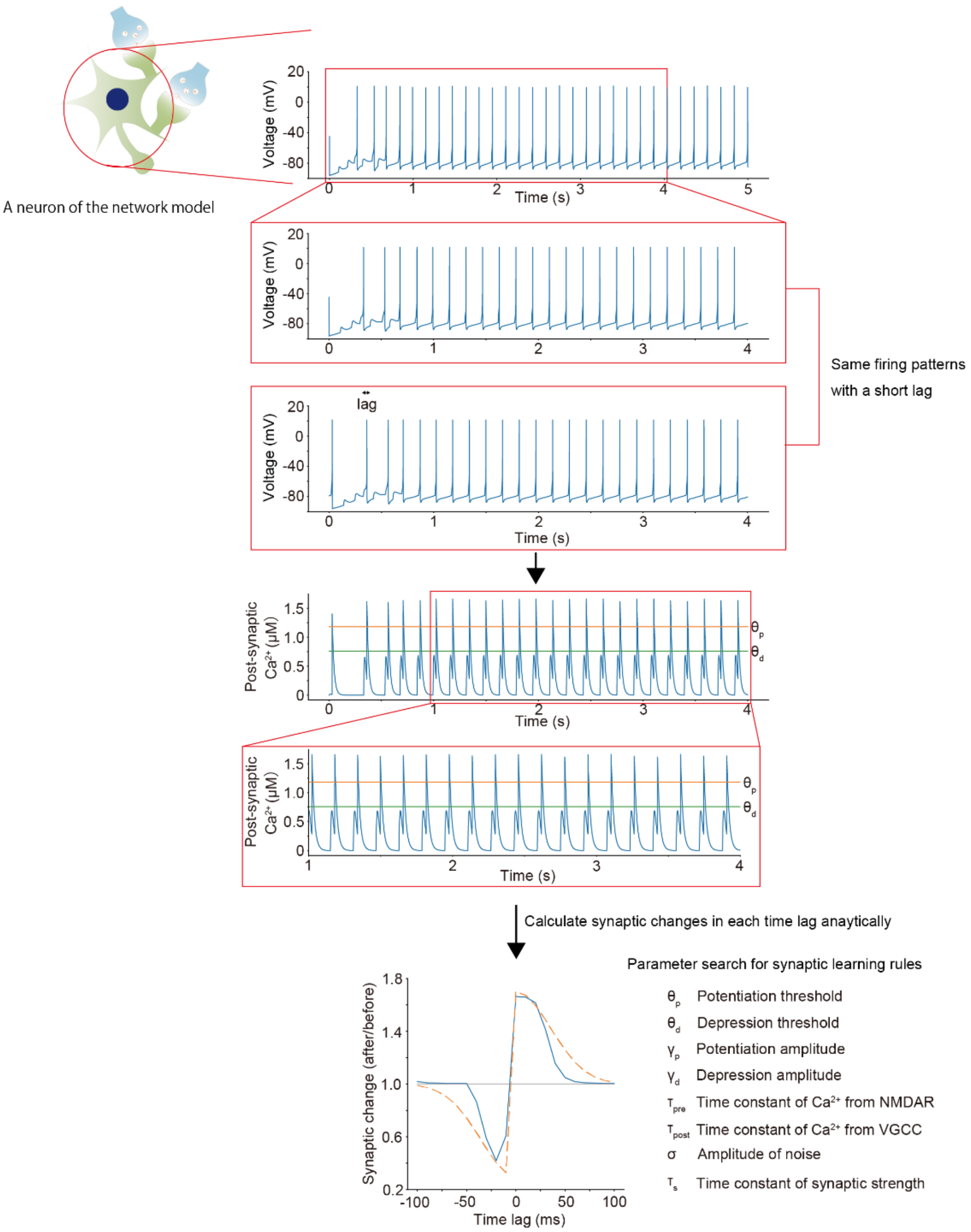


**Fig. S6. Procedures of searching parameter sets for synaptic learning rules in Hodgkin-Huxley-based network model, related to Fig. 3**

The details are shown in *Parameter search for synaptic learning rules in Hodgkin-Huxley-based network models*. A firing pattern of a single neuron was duplicated with a short lag. Then, post-synaptic $Ca^{2+}$ concentrations and synaptic changes in each time lag were analytically calculated in generated parameter sets for synaptic learning rules.

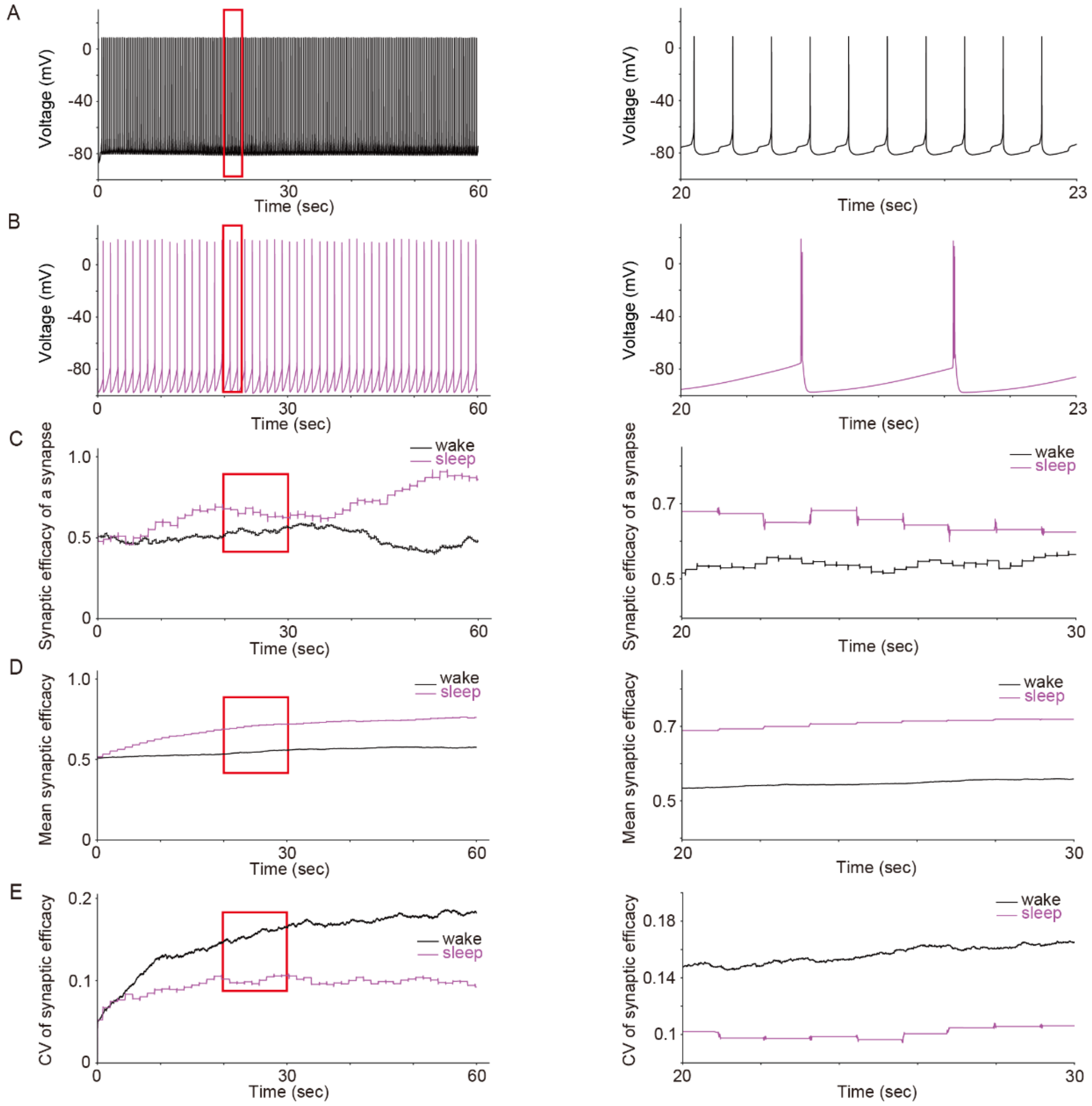


**Fig. S7. Time changes in membrane potential and synaptic efficacy of a representative network model under STDP, related to Fig. 3**

(A-E) Time changes in membrane potential and synaptic efficacy were calculated in a representative intracellular bifurcation model. The parameter set for the channel and receptor conductance of network model was the same as in **Fig. 3C** and the parameter set for STDP learning rule is shown in **Table S8**. Time changes in membrane potential during wake-like patterns (A), membrane potential in sleep-like patterns (B), synaptic efficacy of a representative synapse (C), mean synaptic efficacy (D), and CV of synaptic efficacy (E) are shown. Right figures show enlarged graphs for red rectangles of left figures.

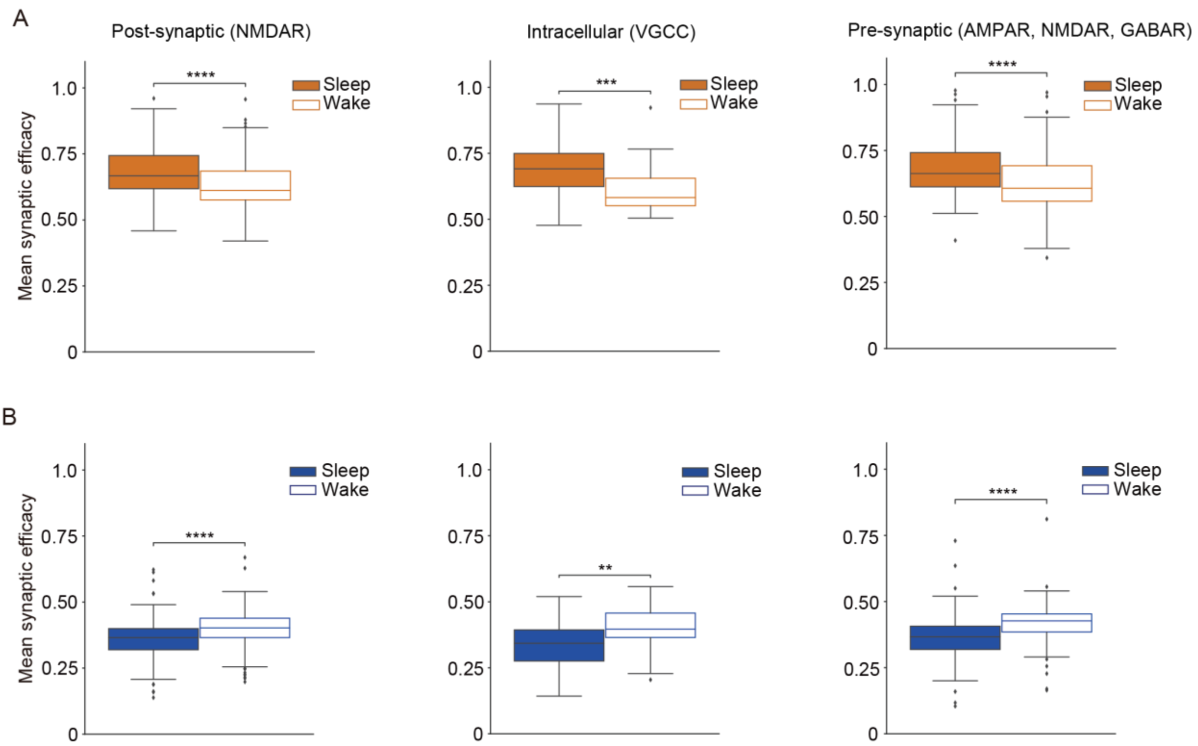


**Fig. S8. Mean synaptic efficacy in Hodgkin-Huxley-based network models under Hebbian and Anti-Hebbian, related to Fig. 3**

(A and B) Box plots for mean synaptic efficacy in sleep-like and wake-like firing patterns under Hebbian (A) and Anti-Hebbian (B) by three types of network models (*n* = 192, 42 and 151 for Hebbian and *n* = 143, 38 and 123 for Anti-Hebbian in post-synaptic, intracellular, and pre-synaptic bifurcation models respectively, *n* represents the number of parameter sets for the network models). The parameter sets for channel or receptor conductance of network models were the same as in **Fig. 3** ***E*** and ***F***. The whiskers above and below of box plots show minimal to maximal values. The box extends from the 25th to the 75th percentile and the middle line indicates the median. * $p < 0.05$, ** $p < 0.01$, *** $p < 0.001$, **** $p < 0.0001$, Student's t-test was applied.

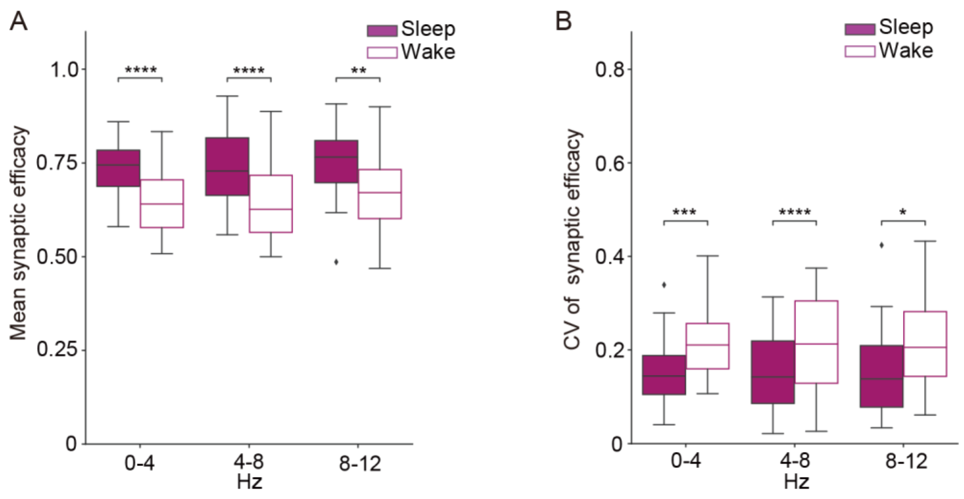


**Fig. S9. Mean synaptic efficacy between sleep-like and wake-like firing patterns at different mean firing rates in Hodgkin-Huxley-based network models, related to Fig. 3**

(A and B) Mean (A) and CV (B) of synaptic efficacy in the post-synaptic bifurcation models under STDP were compared between sleep-like and wake-like firing patterns by different mean firing rates. Models were classified into three groups (0-4 Hz, 4-8 Hz and 8-12 Hz) by mean firing rates during wake-like firing patterns ($n$ = 35, 106 and 29 for each firing rate group, $n$ represents the number of parameter sets for the network models). The box extends from the 25th to the 75th percentile and the middle line indicates the median. * $p < 0.05$, ** $p < 0.01$, *** $p < 0.001$, **** $p < 0.0001$, Student's t-test was applied.

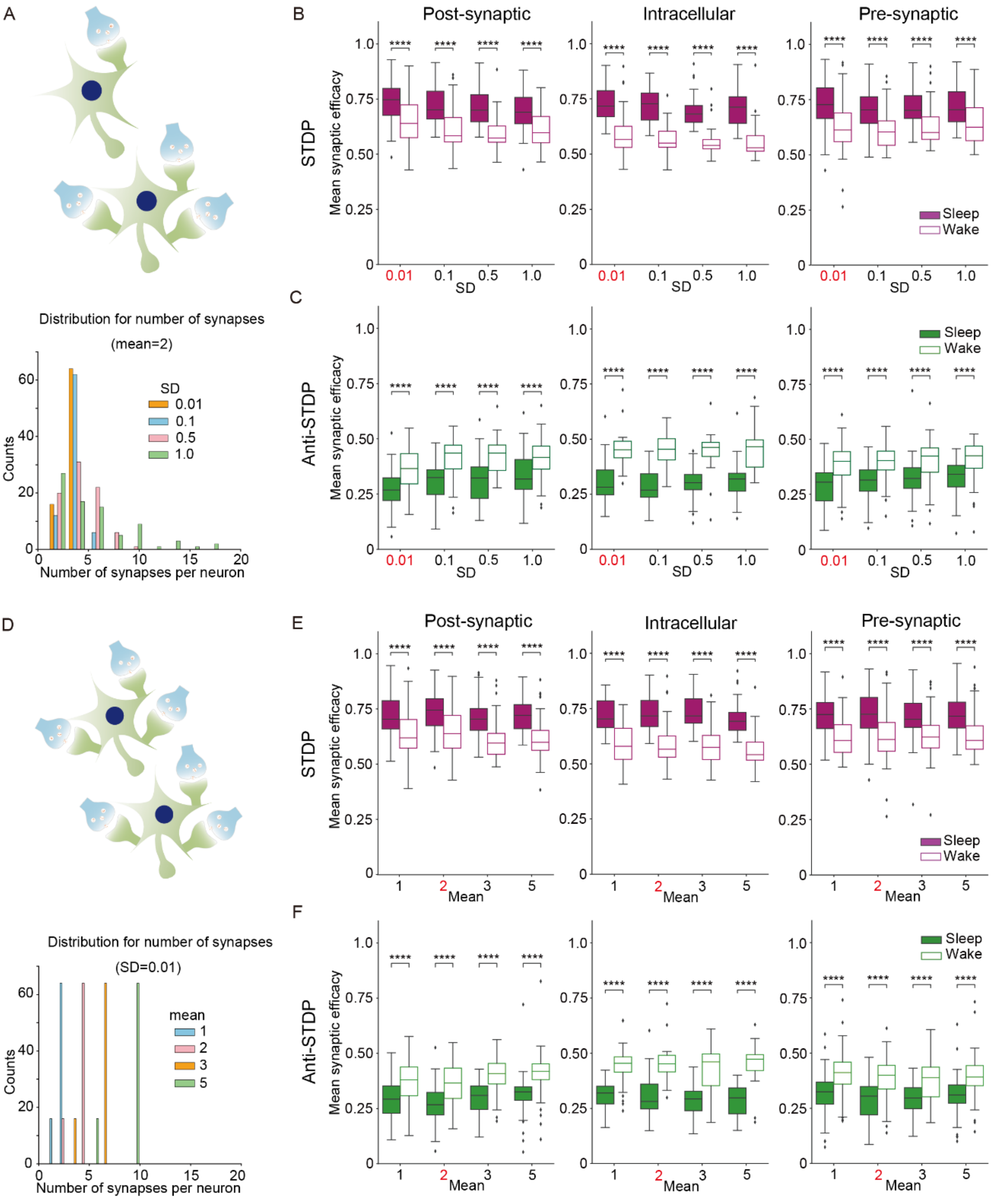


Original

**Fig. S10. Synaptic changes under STDP and Anti-STDP in Hodgkin-Huxley-based network models with different connections, related to Fig. 3**

(A) Schematic illustration and histogram for lognormal distributions of the number of synapses per neuron in different SDs.

(B and C) Box plots for mean synaptic efficacy during sleep-like and wake-like firing patterns under STDP (*n* = 263, 144, 126, 95, *n* = 160, 85, 72, 63, and *n* = 201, 148, 113, 100 for each value of SD in post-synaptic, intracellular and pre-synaptic bifurcation models, respectively) (B) and Anti-STDP (*n* = 121, 82, 57, 49, *n* = 36, 34, 34, 31, and *n* = 119, 90, 61, 58 for each value of SD in post-synaptic, intracellular and pre-synaptic bifurcation models, respectively) (C) in different SDs for lognormal distributions.

(D) Schematic illustration and histogram for lognormal distributions of the number of synapses per neuron in different means.

(E and F) Box plots for mean synaptic efficacy during sleep-like and wake-like firing patterns under STDP (*n* = 135, 263, 149, 146, *n* = 77, 160, 86, 85, and *n* = 147, 201, 142, 145 for each value of mean in post-synaptic, intracellular and pre-synaptic bifurcation models, respectively) (E) and Anti-STDP (*n* = 83, 121, 85, 77, *n* = 36, 36, 35, 38, and *n* = 101, 119, 93, 91 for each value of mean in post-synaptic, intracellular and pre-synaptic bifurcation models, respectively. ) (F) in different means for lognormal distributions.

(B, C, E and F) The whiskers above and below of box plots show minimal to maximal values. The box extends from the 25th to the 75th percentile and the middle line indicates the median. * $p < 0.05$, ** $p < 0.01$, *** $p < 0.001$, **** $p < 0.0001$, Student's t-test was applied. The parameter sets for channel and receptor conductance of network models and synaptic learning rules were the same as in **Fig 3 *E*** and ***F***. *n* represents the number of parameter sets for the network models.

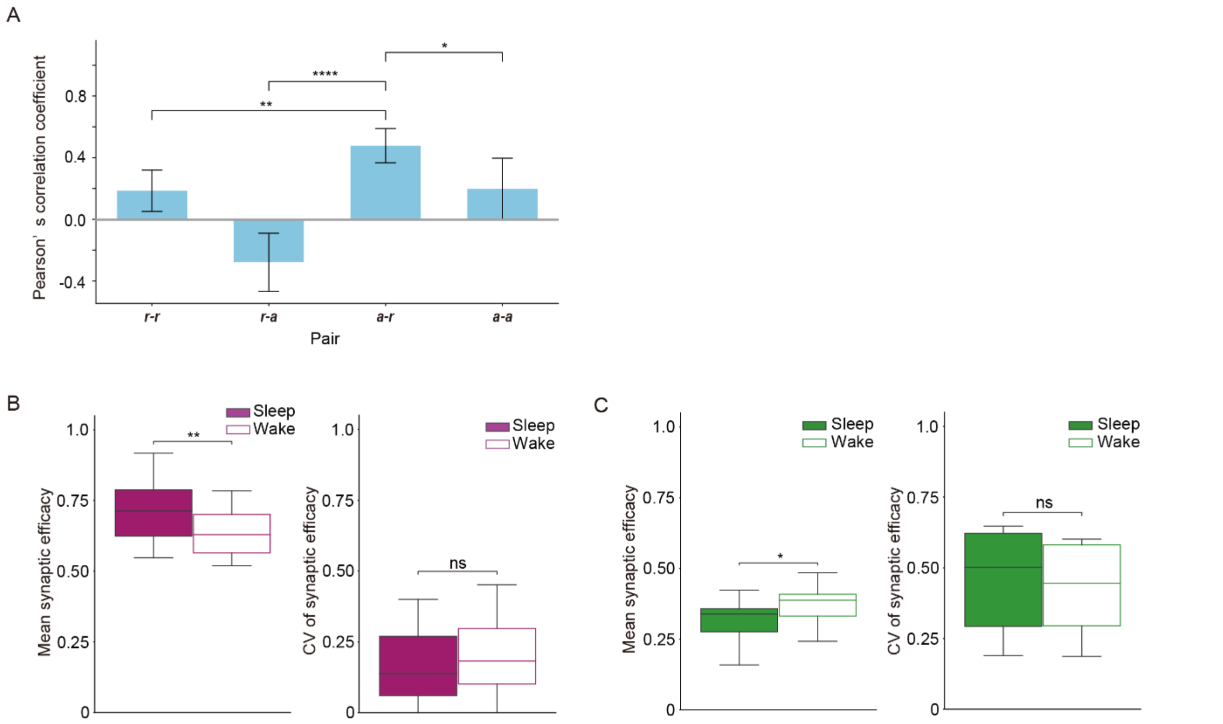


**Fig. S11. Analysis in multiple parameter sets for Hodgkin-Huxley-based network models bifurcated by the post-synaptic mechanism under STDP or Anti-STDP with sleep-wake dynamics, related to Fig. 4**

(A) Pearson's correlation coefficients were calculated in combinations of ***r*** and ***a*** with multiple parameter sets for Hodgkin-Huxley-based network models. NMDAR conductance was updated by ***r*** or ***a*** and parameters for sleep-wake dynamics were optimized by Pearson's correlation coefficients between Process S and ***r*** or ***a*** (For example, ***r-a*** means that conductance was updated by ***r*** and parameters for sleep-wake dynamics were optimized by Pearson's correlation coefficients between Process S and ***a***). The network structures and initial values for variables (***a***, ***r*** and $\xi$) were the same as in **Fig. 4**. The simulations were conducted for 300 seconds. *n* = 14, 14, 15, and 15 for ***r-r***, ***r-a***, ***a-r***, and ***a-a*** pair, respectively. * $p < 0.05$, ** $p < 0.01$, *** $p < 0.001$, **** $p < 0.0001$, Student's t-test was applied.

(B) Box plots for mean and CV of synaptic efficacy during sleep-like and wake-like periods in multiple parameter sets for network models with sleep-wake dynamics under STDP (*n* = 26) and Anti-STDP (*n*=12). Initial values for variables (***a***, ***r*** and $\xi$) were the same as in the representative models in **Fig. 4**.

NMDAR conductance was updated by ***a*** and optimized by Pearson's correlation coefficients between Process S and ***r***. The whiskers above and below of box plots show minimal to maximal values. The box extends from the 25th to the 75th percentile and the middle line indicates the median. * $p < 0.05$, ** $p < 0.01$, *** $p < 0.001$, **** $p < 0.0001$, Student's t-test was applied.

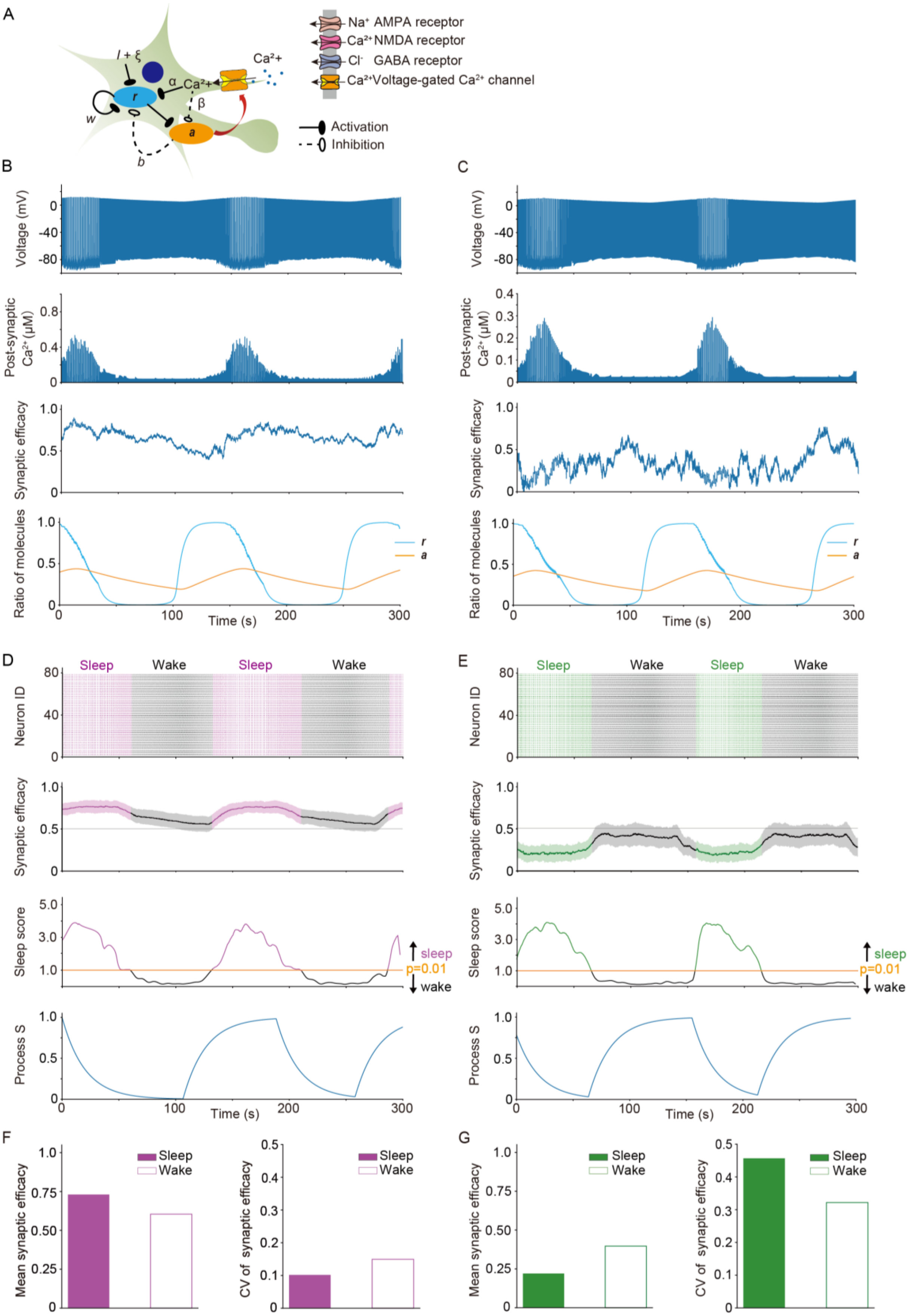
A
Na⁺ AMPA receptor
Ca²⁺ NMDA receptor
Cl⁻ GABA receptor
Ca²⁺ Voltage-gated Ca²⁺ channel
Activation
Inhibition
I + ξ
Ca²+
α
β
w
r
a
b
B
C
Voltage (mV)
Post-synaptic Ca²⁺ (μM)
Synaptic efficacy
Ratio of molecules
r
a
Time (s)
D
E
Sleep
Wake
Neuron ID
Sleep score
sleep
p=0.01
wake
Process S
F
G
Mean synaptic efficacy
CV of synaptic efficacy
Sleep
Wake

**Fig. S12. Representative models with sleep-wake dynamics under STDP and Anti-STDP in the intracellular bifurcation model, related to Fig. 4**

VGCC conductance was updated by ***a*** and parameters for the sleep-wake dynamics model were optimized by Pearson's correlation coefficient between Process S and ***r***. The parameter set for channel or receptor conductance of network models, synaptic learning rules and sleep-wake dynamics and initial values for variables in a representative model are shown in **Tables S5-S8**. The simulations were conducted for 500 seconds.

(A) Schematic illustration of the model for sleep-wake dynamics in the intracellular bifurcation mechanism. ***r***, ***a***, $\alpha$, $\beta$, *w*, and *b* corresponds to variables defined in the equations of the sleep-wake dynamics (*Materials and Methods* in the main text).

(B and C) Time changes in membrane potential of a neuron and post-synaptic $Ca^{2+}$, synaptic efficacy and ratio of two phosphorylated states of kinases (***r*** and ***a***) of a synapse in representative network models under STDP (B) and Anti-STDP (C). The results from 200-500 seconds are shown.

(D and E) Raster plots and time changes in mean synaptic efficacy, sleep score and Process S in representative network models under STDP (D) and Anti-STDP (E). The shadow in time changes in mean synaptic efficacy represents SD. The network was considered to be in the sleep-like or wake-like states if the sleep score was above or below the threshold, respectively (the threshold is the value of sleep score where p = 0.01, see *Evaluation of synchronization and desynchronization in Hodgkin-Huxley-based network models*). The results of 200-500 seconds are shown.

(F and G) Mean and CV of synaptic efficacy during the periods of sleep-like and wake-like states in representative network models under STDP (F) and Anti-STDP (G).

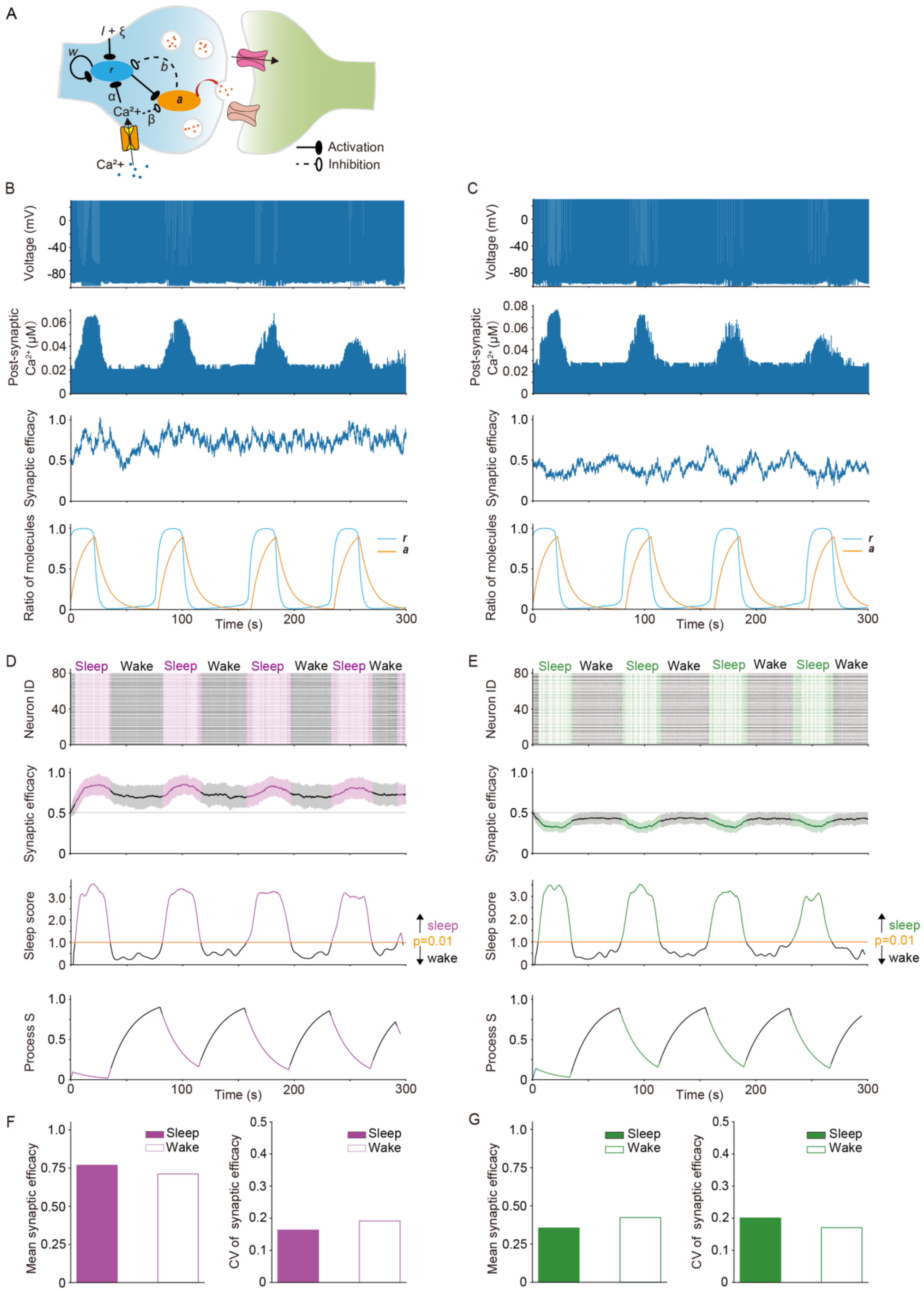
A
I + ξ
w
r
b
a
α
β
Ca²+
Ca²+
Activation
Inhibition
B
C
Voltage (mV)
Post-synaptic Ca²⁺ (μM)
Synaptic efficacy
Ratio of molecules
r
a
Time (s)
D
E
Sleep
Wake
Neuron ID
Synaptic efficacy
Sleep score
sleep
p=0.01
wake
Process S
Time (s)
F
G
Mean synaptic efficacy
CV of synaptic efficacy
Sleep
Wake

**Fig. S13. Representative models with sleep-wake dynamics under STDP and Anti-STDP learning rules in the pre-synaptic bifurcation model, related to Fig. 4**

Coefficients for pre-synaptic activations were updated by ***a*** and parameters for the sleep-wake dynamics model were optimized by Pearson's correlation coefficient between Process S and ***r***. The parameter set for channel or receptor conduces of network models, synaptic learning rules and sleep-wake dynamics and initial values for variables in a representative model are shown in **Tables S5-S8**. The simulations were conducted for 300 seconds.

(A) Schematic illustration of the model for sleep-wake dynamics in the intracellular bifurcation mechanism. ***r***, ***a***, $\alpha$, $\beta$, *w*, and *b* corresponds to variables defined in the equations of the sleep-wake dynamics (*Materials and Methods* in the main text).

(B and C) Time changes in membrane potential of a neuron and post-synaptic $Ca^{2+}$, synaptic efficacy and ratio of two phosphorylated states of kinases (***r*** and ***a***) of a synapse in representative network models under STDP (B) and Anti-STDP (C).

(D and E) Raster plots and time changes in mean synaptic efficacy, sleep score and Process S in representative network models under STDP (D) and Anti-STDP (E). The shadow in time changes in mean synaptic efficacy represents SD. The network was considered to be in the sleep-like or wake-like states if the sleep score was above or below the threshold, respectively (the threshold is the value of sleep score where p = 0.01, see *Evaluation of synchronization and desynchronization in Hodgkin-Huxley-based network models*).

(F and G) Mean and CV of synaptic efficacy during the periods of sleep-like and wake-like states in representative network models under STDP (F) and Anti-STDP (G).

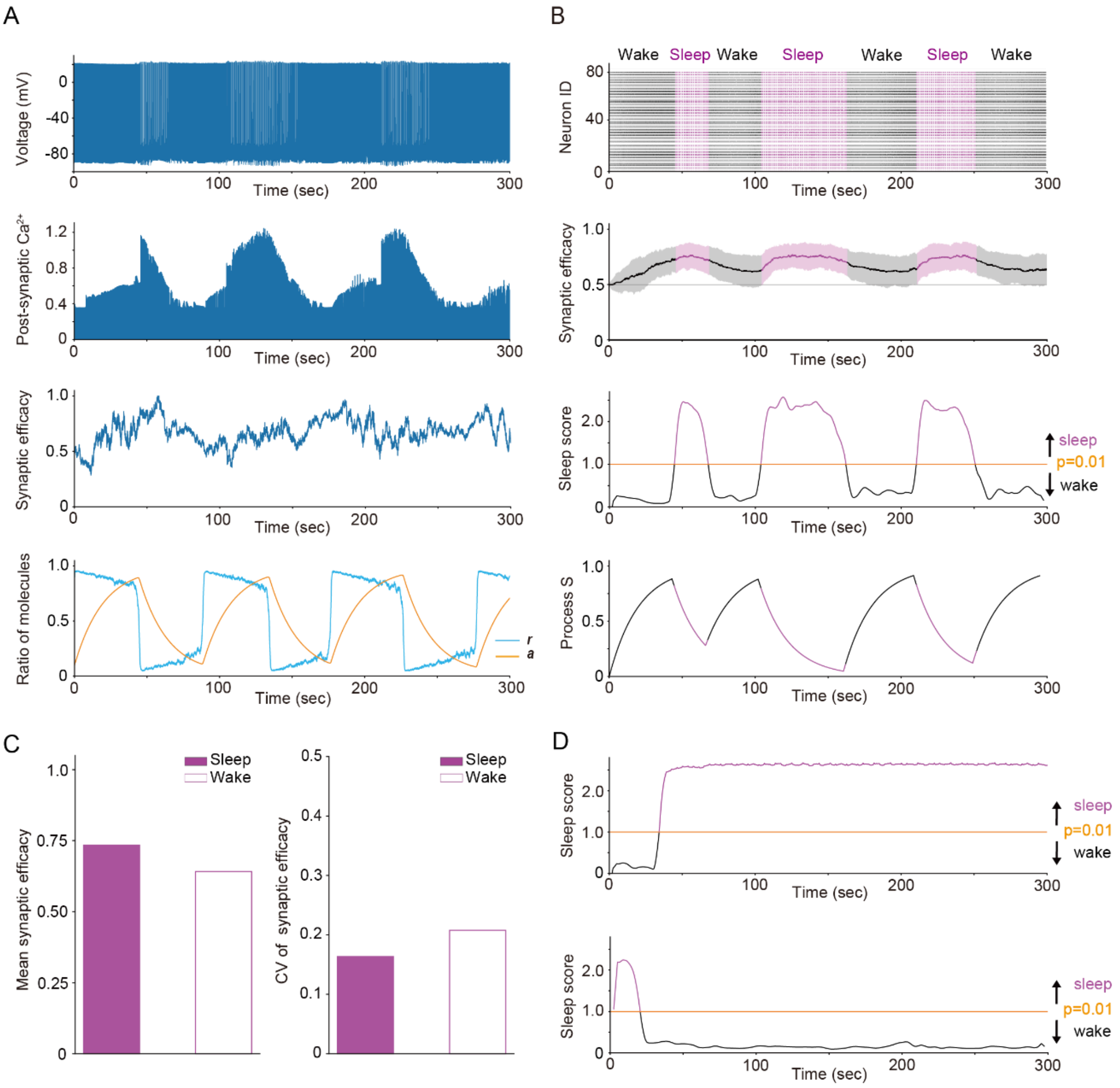


**Fig. S14. Sleep-wake dynamics in the bistable regime, related to Fig. 4**

The model for sleep-wake dynamics can represent multiple regimes. In the oscillatory regime (**Fig. 4**), activity alternates between sleep-like and wake-like states. In the bistable regime, although sleep-like and wake-like states are relatively stable, sufficiently large noises induces alternations between two states, resulting in variable durations of two states. Synaptic efficacy was calculated in a representative network model bifurcated by the post-synaptic mechanism with sleep-wake dynamics and STDP learning rule in the bistable regime. The conductance of NMDAR was updated by ***a*** and the simulations were optimized by Pearson's correlation coefficients between Process S and ***r***. The simulations were conducted for 300 seconds. The parameter set for channel or receptor conduces of network models, synaptic learning rules and sleep-wake dynamics and initial values for variables in a representative model are shown in **Tables S5-S8**.

(A) Time changes in membrane potentials, post-synaptic $Ca^{2+}$, synaptic efficacy, and ratio of two phosphorylated states of kinases (***r*** and ***a***) in a single neuron of a representative network model.

(B) Raster plots, time changes in mean synaptic efficacy, sleep score, and Process S in a representative network model. The shadow in time changes in mean synaptic efficacy represents SD. The network was considered to be in the sleep-like or wake-like states if the sleep score was above or below the threshold, respectively (the threshold is the value of sleep score where p = 0.01, see *Evaluation of synchronization and desynchronization in Hodgkin-Huxley-based network models*)

(C) Mean and CV of synaptic efficacy during sleep-like and wake-like periods in a representative network model.

(D) The results of simulations without noise. Simulations were conducted in $\theta = 0$ and $\xi = 0$ (the initial value for $\xi$ is also 0) while other parameters were the same as in simulations with noise. The upper graph shows the time change in sleep scores in the simulation starting from wake-like firing patterns and the lower graph shows the time change in sleep scores starting from the sleep-like firing patterns.

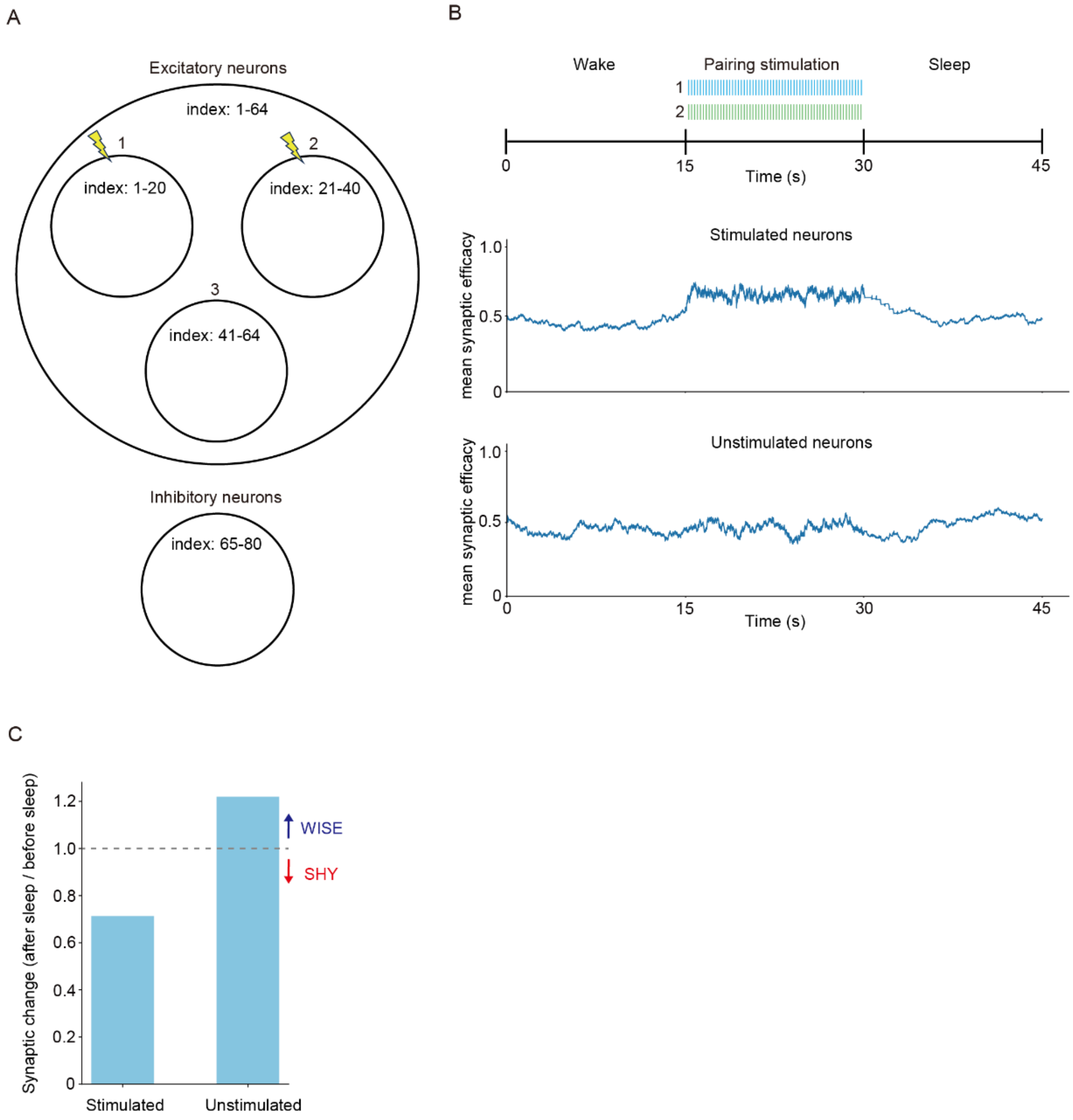


**Fig. S15. Synaptic changes in Hodgkin-Huxley-based network models with stimulation during wakefulness under STDP, related to Fig. 5**

Synaptic efficacy was calculated in a representative network model bifurcated by the intracellular mechanism under STDP. Parameter sets for the channel or receptor conductance and the synaptic learning rule are shown in **Tables S5** and **S8** respectively. The VGCC conductance was multiplied by $10^{-0.4}$ and $10^{-0.1}$ to its original value to generate wake-like and sleep-like firing patterns. Stimulations were applied for 15 seconds at 20 Hz after wake-like firing patterns. The stimulation was optimized by changing its waveforms and rates so that the potentiation of synaptic efficacy between stimulated groups were observed (see *Calculation of synaptic efficacy under synaptic learning rules in Hodgkin–Huxley-based network models including stimulation during the wakefulness*).

(A) Schematic illustration for grouping excitatory neurons. Group **1** and **2** were stimulated.

(B) Time changes in mean synaptic efficacy of stimulated neurons and unstimulated neurons.

(C) Ratio of mean synaptic efficacy after and before sleep-like firing patterns in stimulated and unstimulated neurons.

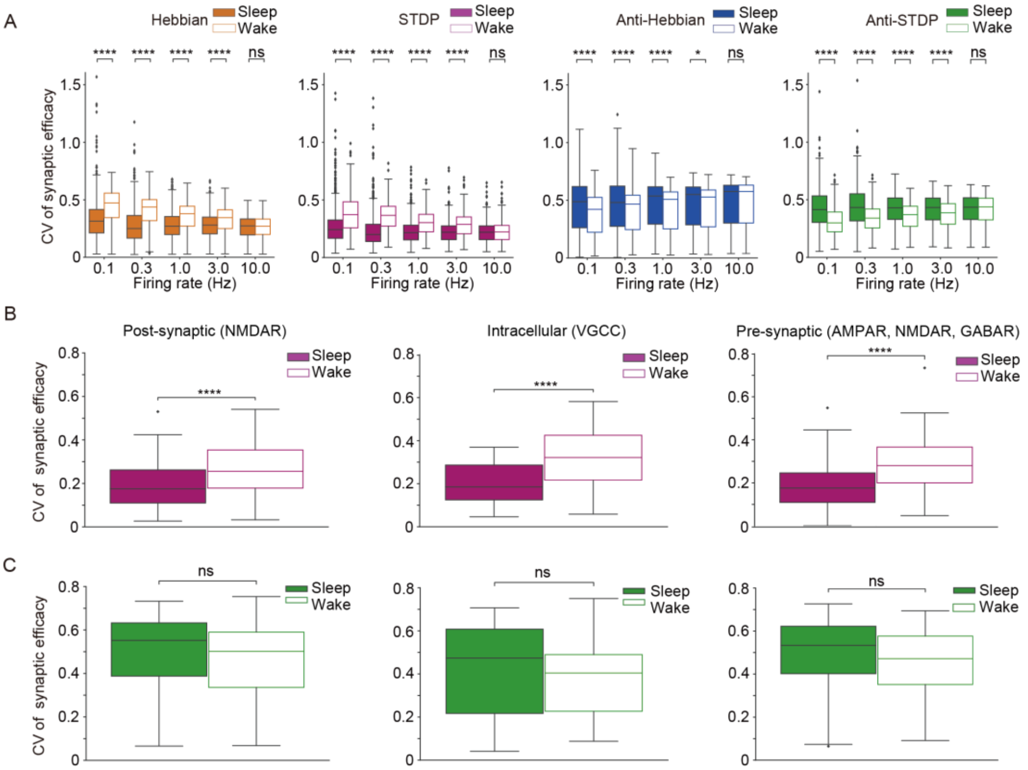


**Fig. S16. CV of synaptic efficacy under different synaptic learning rules**

(A) Box plots for CV of synaptic efficacy in sleep-like and wake-like firing patterns by synaptic learning rules and mean firing rates (*n* = 1000 for each firing rate, *n* represents the number of synaptic learning rules). The network structure, firing patterns and parameters for synaptic learning rules are the same as in **Figure 1H**.

(B and C) Box plots for CV of synaptic efficacy during sleep-like and wake-like firing patterns under STDP (B) and Anti-STDP (C) in Hodgkin-Huxley-based network models (*n* = 191, 52 and 150 for STDP and *n* = 121, 36 and 119 for Anti-STDP in post-synaptic, intracellular, and pre-synaptic bifurcation models respectively. *n* represents the number of parameter sets for the network models).

(A-C) The whiskers above and below of box plots show minimal to maximal values. The box extends from the 25th to the 75th percentile and the middle line indicates the median. * $p < 0.05$, ** $p < 0.01$, *** $p < 0.001$, **** $p < 0.0001$, Welch's t-test was applied in (A) and Student's t-test was applied in (B) and (C).

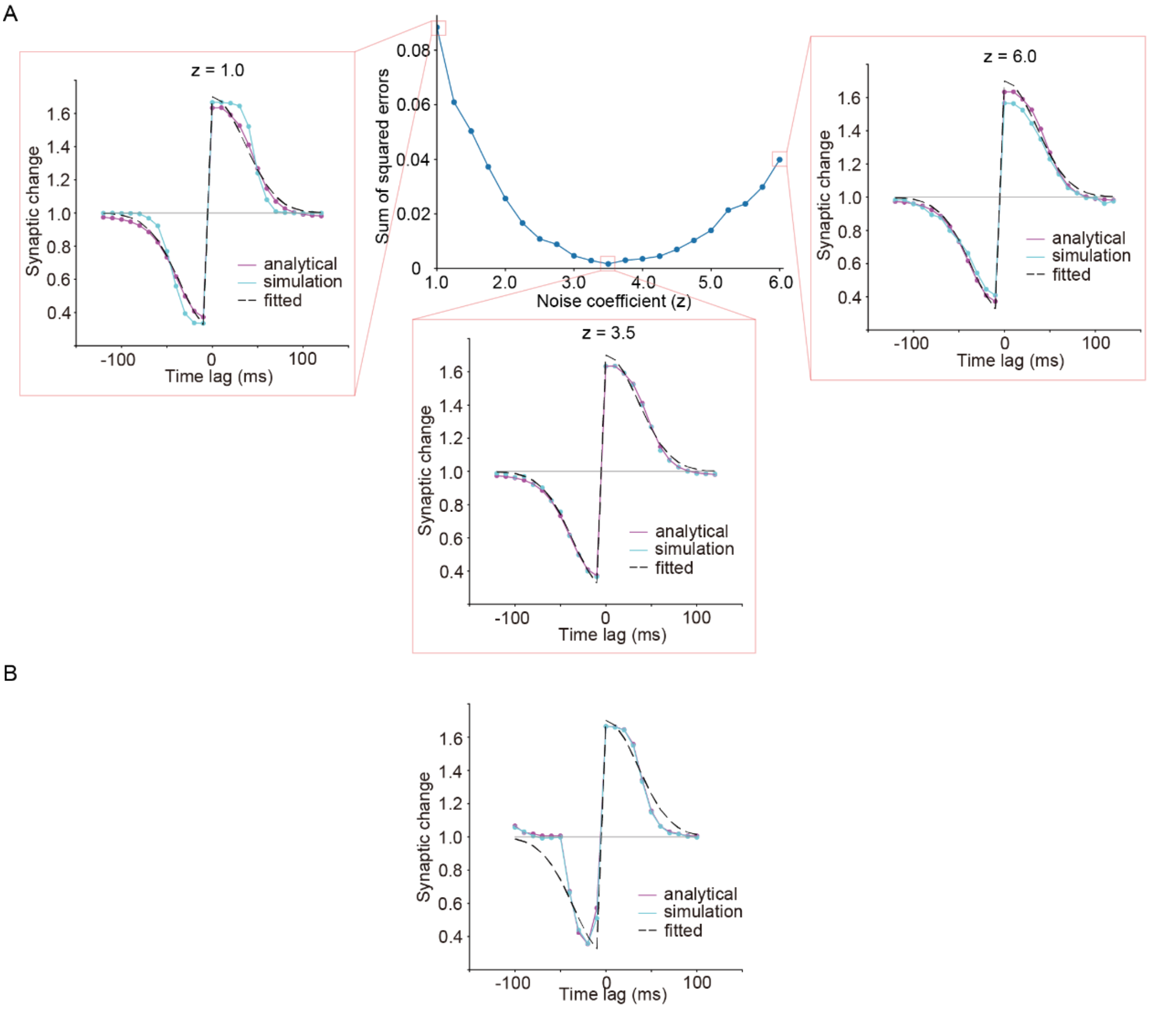


**Fig. S17. Coefficient of noise and comparison of analytical and simulation results**

(A) The sum of squared errors (SSE) between analytical solutions and simulation results were calculated in different noise coefficient (a center figure). All simulations were conducted in step size = 0.1. The simulation results were compared with analytical solutions and fitting curves (surrounding figures). The parameter set for STDP learning rule is shown in **Table S8**.

(B) Comparison of analytical and simulation results in parameter search for STDP learning rule in a representative network model bifurcated by the post-synaptic mechanism. The parameter set for channel or receptor conductance of network model was the same as in **Fig. 3 *C*** and the parameter set for STDP learning rule is shown in **Table S8**. The simulation was conducted by step size = 0.1.

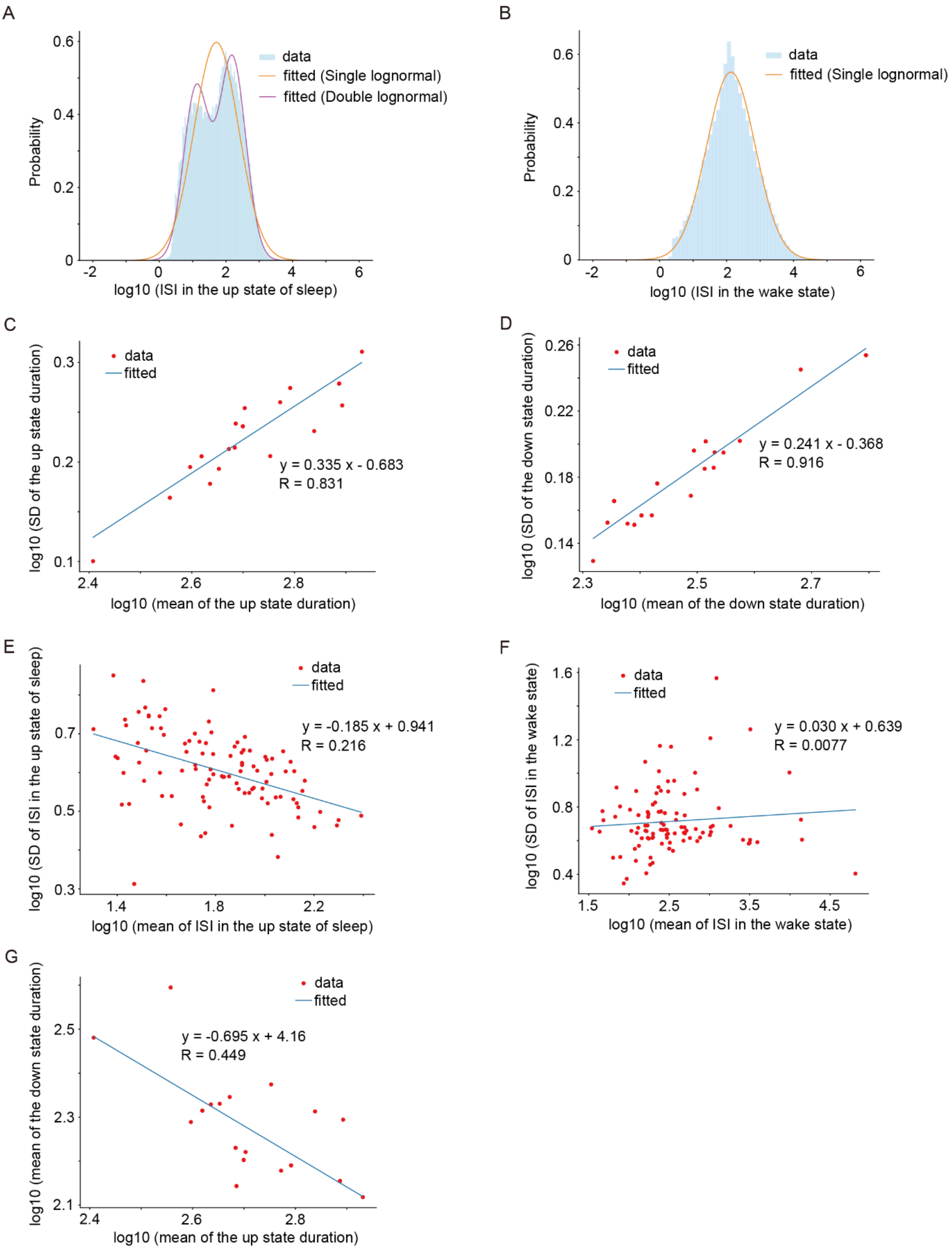
A
data
fitted (Single lognormal)
fitted (Double lognormal)
Probability
log10 (ISI in the up state of sleep)
B
data
fitted (Single lognormal)
Probability
log10 (ISI in the wake state)
C
data
fitted
y = 0.335 x - 0.683
R = 0.831
log10 (SD of the up state duration)
log10 (mean of the up state duration)
D
data
fitted
y = 0.241 x - 0.368
R = 0.916
log10 (SD of the down state duration)
log10 (mean of the down state duration)
E
data
fitted
y = -0.185 x + 0.941
R = 0.216
log10 (SD of ISI in the up state of sleep)
log10 (mean of ISI in the up state of sleep)
F
data
fitted
y = 0.030 x + 0.639
R = 0.0077
log10 (SD of ISI in the wake state)
log10 (mean of ISI in the wake state)
G
data
fitted
y = -0.695 x + 4.16
R = 0.449
log10 (mean of the down state duration)
log10 (mean of the up state duration)

**Fig. S18. Distributions of ISI and linear regression analysis for spike trains in vivo data**

(A and B) We investigated the distribution for ISI of the spike-train data for all excitatory neurons (verified by cross-correlogram) in a dataset of a previous article (3, 4). Distributions for ISI in Up states in the state of sleep (A) and wake (B) are shown. The lognormal distributions fitted well in the ISI of the state of wake (B). Although mixed lognormal distributions were expected in the ISI of the sleep Up states, we assumed a single lognormal distribution in simulations for simplification (A).

(C-G) We performed the linear regression analysis on the mean and SD of Up-state duration (C), the mean and SD of Down-state duration (D), the mean and SD of ISI in the state of wake (E) and in the Up states of sleep (F) and the mean Up-state duration and mean Down-state duration (G).

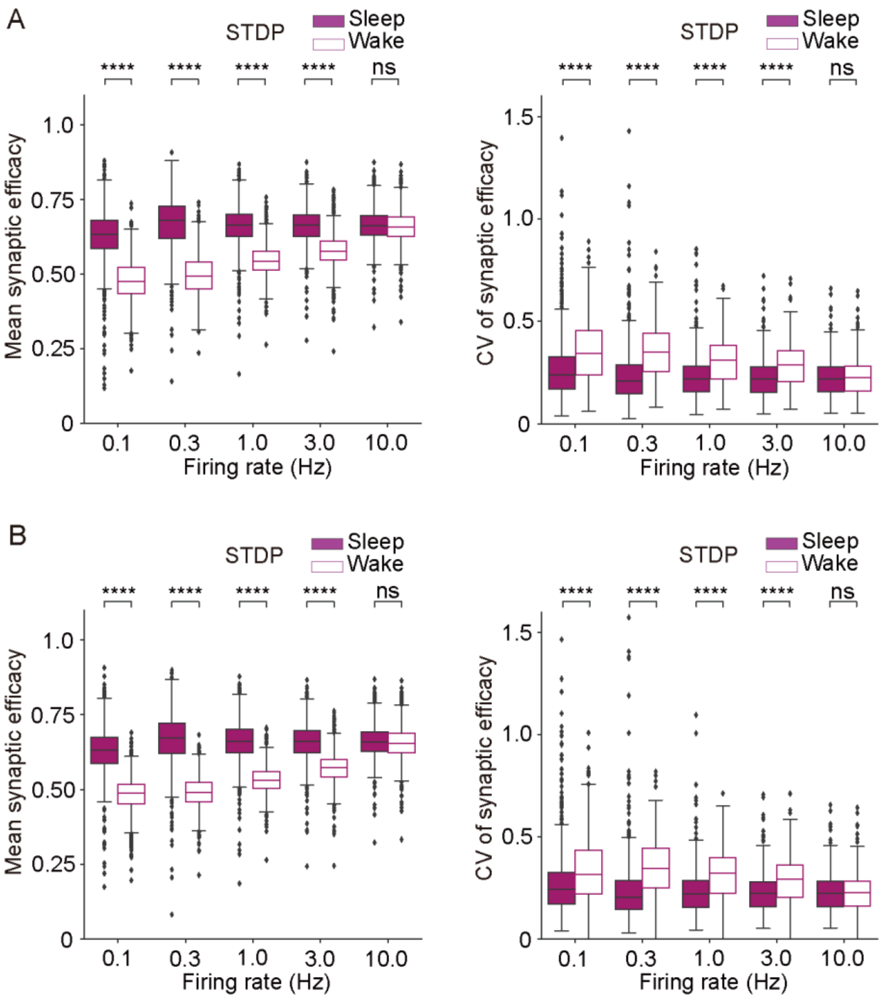


**Fig. S19. Mean synaptic efficacy in different membrane potential differences between Up and Down states.**

(A and B) Box plots for mean synaptic efficacy during sleep-like and wake-like firing patterns with 10mV (A) and 5mV (B) membrane potential differences are shown ($n$ = 1000 for each firing rate, n represents the number of synaptic learning rules). The parameter sets for STDP and spike trains are the same as in **Fig. 1*H***. The parameters for constructing waveforms are shown in **Table S3**. The whiskers above and below of box plots show minimal to maximal values. The box extends from the 25th to the 75th percentile and the middle line indicates the median. * $p < 0.05$, ** $p < 0.01$, *** $p < 0.001$, **** $p < 0.0001$, Welch's t-test was applied.

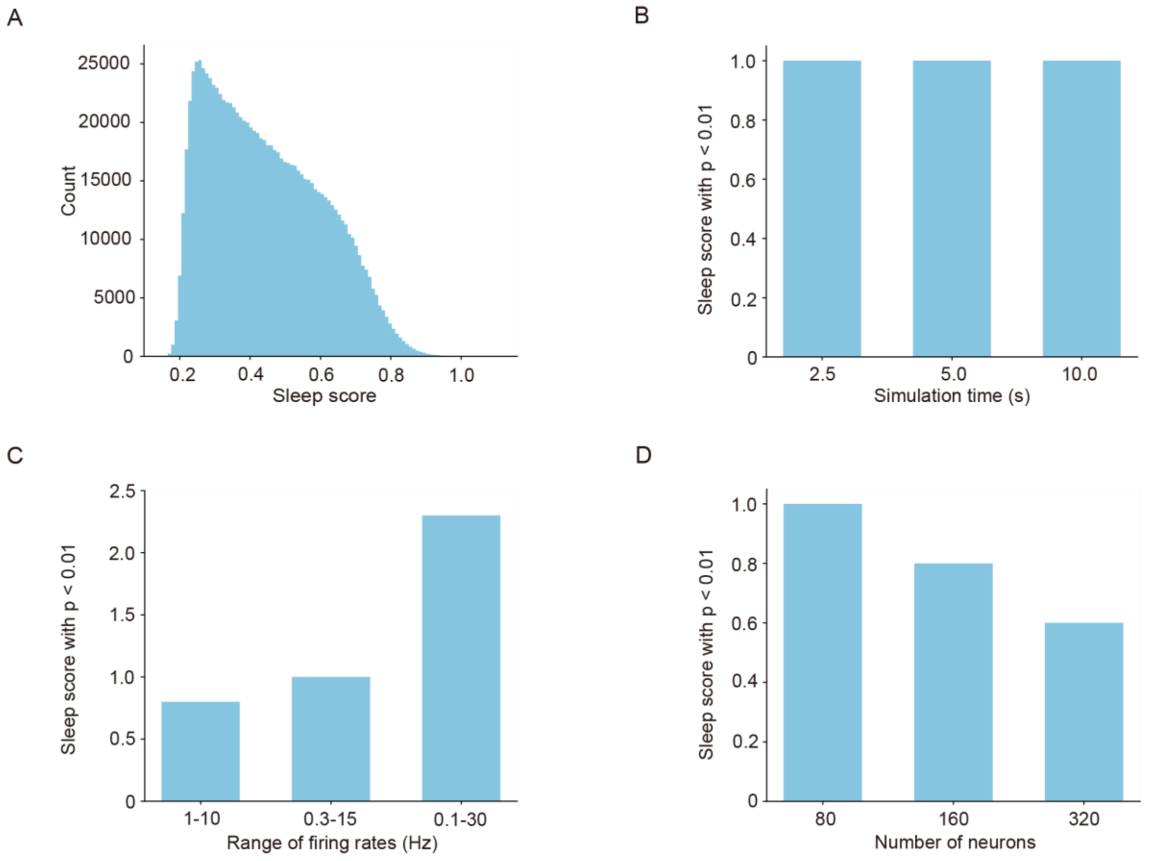


**Fig. S20. Distribution of sleep scores and sleep scores with $p < 0.01$ in other conditions**

(A) Distribution of sleep scores in 1,000,000 desynchronized spikes in the condition of 80 neurons, 5 seconds' simulation time and 0.5-15 Hz. Wake-like desynchronized spikes were sampled from lognormal distributions for ISI (see *Definition of lognormal distributions based on in vivo recordings*). (B-D) sleep scores with $p < 0.01$ in different simulation times (B), ranges of mean firing rates (C) and number of neurons (D). P values were calculated by the distribution of 1,000,000 desynchronized spikes of each condition.

#### Supplementary Tables

**Table S1. Fixed values in simple model for synaptic learning rules**

Values were based on the previous study (8) and used in **Figs. 1, 2** and **5*A*,** and **Figs. S2, S3, S16*A*, S17*A*** and **S19**.

| | | | |
|---|---|---|---|
| $A$ | Area of neuron | 0.02 | $mm^2$ |
| $V_{Ca}$ | $Ca^{2+}$ reversal potential | 120 | mV |
| $V_{rest}$ | Resting potential | -70 | mV |
| $\alpha_{Ca}$ | Coefficient of $Ca^{2+}$-entry | 0.5 | μM/(nA ms) |
| $a_{sNMDA}$ | Coefficient of $s_{NMDA}$ | 5 | |
| $\tau_{sNMDA}$ | Time constant of $s_{NMDA}$ | 10 | ms |
| $a_{xNMDA}$ | Coefficient of $x_{NMDA}$ | 34.8 | |
| $\tau_{xNMDA}$ | Time constant of $x_{NMDA}$ | 0.2 | ms |
| $g_{NMDA}$ | The conductance of the NMDA receptors | 0.0138403 | μS |
| $g_{Ca\_spine}$ | The conductance of the voltage-gated $Ca^{2+}$ channels in spines | 0.050580 | $mS/cm^2$ |

**Table S2. Value ranges of parameters for synaptic learning rules**

Parameter values were randomly sampled from uniform distributions within these ranges of values when searching the parameter sets for synaptic learning rules fitting gaussian curves (**Figs. 1 *E*** and ***F,*** and **Fig. S1**).

| **Parameter** | **min** | **max** | **unit** |
|---|---|---|---|
| $\theta_p$ | 0.06 | 1.6 | |
| $\theta_d$ | 0.06 | 1.6 | |
| $\gamma_p$ | 50 | 5000 | |
| $\gamma_d$ | 50 | 10000 | |
| $\tau_{pre}$ | 1 | 100 | µM |
| $\tau_{post}$ | 1 | 100 | µM |
| $\sigma$ | 0.5 | 35 | |
| $\tau_s$ | 2500 | 2500000 | ms |

**Table S3. Parameters for constructing voltage waveforms from spike trains**

These are values used in **Figs. 1, 2** and **5*A***, and **Figs. S2, S3, S16*A*** and **S17*A*** to construct voltage waveforms from spike trains.

| **Parameter** | **Figs. 1, 2, 5*A*, S2, S3, S16*A*,** and **S17*A*** | **Fig. S19*A*** | **Fig. S19*B*** |
|---|---|---|---|
| peak of action potential (mV) | 13 | 13 | 13 |
| Up state (mV) | 61 | 66 | 71 |
| Down state (mV) | 76 | 76 | 76 |
| duration of action potential (ms) | 1.6 | 1.6 | 1.6 |

**Table S4. Fixed values in the Hodgkin-Huxley-based network model**

Values were based on the previous study (8) and used in simulations of Hodgkin-Huxley-based network models.

| | | | |
|---|---|---|---|
| $C$ | Membrane capacitance | 1 | μF/cm2 |
| $A$ | Area of neuron | 0.02 | mm2 * |
| $V_L$ | Leak reversal potential | -60.95 | mV |
| $V_{Na}$ | Sodium reversal potential | 55 | mV |
| $V_K$ | Potassium reversal potential | -100 | mV |
| $\tau_{hA}$ | Time-constant of *h*A | 15 | ms |
| $V_{Ca}$ | $Ca^{2+}$ reversal potential | 120 | mV |
| $K_D$ | Dissociation constant of $Ca^{2+}$-dependent K+ channels | 30 | μM |
| $V_{AMPA}$ | APMA receptor reversal potential | 0 | mV |
| $V_{NMDA}$ | NMDA receptor reversal potential | 0 | mV |
| $V_{GABA}$ | GABA receptor reversal potential | -70 | mV |
| $\alpha_{Ca}$ | Coefficient of $Ca^{2+}$-entry | 0.5 | μM/(nA ms) |
| $a_{AMPA}$ | Coefficient of $s_{AMPA}$ | 3.48 | |
| $\tau_{AMPA}$ | Time-constant of $s_{AMPA}$ | 2 | ms |
| $a_{sNMDA}$ | Coefficient of $s_{NMDA}$ | 5 | |
| $\tau_{sNMDA}$ | Time-constant of $s_{NMDA}$ | 10 | ms |
| $a_{xNMDA}$ | Coefficient of $x_{NMDA}$ | 34.8 | |
| $\tau_{xNMDA}$ | Time-constant of $x_{NMDA}$ | 0.2 | ms |
| $a_{GABA}$ | Coefficient of $s_{GABA}$ | 1 | |
| $\tau_{GABA}$ | Time-constant of $s_{GABA}$ | 10 | ms |

**Table S5. The representative parameter sets for Hodgkin-Huxley-based network models**

These parameter sets were used in the network models by three types of bifurcations in **Figs. 3*C*** and **4,** and **Figs. S7**, and **S12-S17**.

| **Parameter** | **Post-synaptic (NMDAR)** | **Intracellular (VGCC)** | **Pre-synaptic (AMPAR, NMDAR, GABAR)** | **Intracellular (VGCC, stimulation)** |
|---|---|---|---|---|
| $g_L$ [mS/cm2] | 0.019148334 | 0.022759897 | 0.01347216 | 0.027854564 |
| $g_{Na}$ [mS/cm2] | 8.874191845 | 1.11512535 | 97.2559829 | 1.52942181 |
| $g_K$ [mS/cm2] | 36.58825101 | 20.17731588 | 21.5320075 | 45.4733042 |
| $g_A$ [mS/cm2] | 0.0537862 | 0.012954267 | 0.07987940 | 0.971993713 |
| $g_{KS}$ [mS/cm2] | 0.970689584 | 0.108535743 | 0.51776950 | 0.063020293 |
| $g_{NaP}$ [mS/cm2] | 0.484254937 | 1.024188842 | 0.16093585 | 1.247766538 |
| $g_{AR}$ [mS/cm2] | 0.011875021 | 0.011250618 | 0.01162880 | 0.026532032 |
| $g_{Ca}$ [mS/cm2] | 0.414856695 | 0.58317768 | 0.96499122 | 1.608910891 |
| $g_{KCa}$ [mS/cm2] | 14.29704464 | 0.882928362 | 7.631158679 | 0.915449325 |
| $g_{AMPA}$ [μS] | 0.018769194 | 0.457834492 | 1.602118176 | 0.148544478 |
| $g_{NMDA}$ [μS] | 0.033924532 | 0.132922379 | 0.001826 | 0.029571348 |
| $g_{GABA}$ [μS] | 0.023480694 | 0.540419577 | 0.00171828 | 1.930015913 |
| $\tau_{Ca}$ [ms] | 347.6255975 | 978.4720314 | 114.020606 | 246.6238776 |

**Table S6. The representative parameter sets for sleep-wake dynamics**

These parameter sets were used in simulations of the Hodgkin-Huxley-based network models with sleep-wake dynamics in **Fig. 4** and **Figs. S12-S14**.

| Parameter | Post-synaptic (Oscillation, STDP) | Post-synaptic (Bistable, STDP) | Intracellular (Oscillation, STDP) | Pre-synaptic (Oscillation, STDP) |
|---|---|---|---|---|
| $\alpha$ | 300.0 | 0.2 | 10.5 | 1100000.0 |
| $\beta$ | 0.0041 | 0.0005 | 0.005 | 0.25 |
| $b$ | 60.0 | 1.0 | 400 | 80.0 |
| $c$ | 0.2 | 1.0 | 0.2 | 0.25 |
| $d$ | 0.8 | 0.5 | 0.7 | 0.65 |
| $e$ | 15.0 | 15.0 | 8.0 | 80.0 |
| $\tau_r$ (ms) | 13000 | 200 | 6000 | 2500 |
| $\tau_a$ (ms) | 60000 | 20000 | 100000 | 10000 |
| $w$ | 70.0 | 6.3 | 100.0 | 100.0 |
| $I$ | -20.0 | -2.65 | -20.0 | -18.0 |
| $\theta$ | 0 | 0.6 | 0 | 0 |
| $\varepsilon$ | 0 | 0.7 | 0 | 0 |
| Max_rate | 8.5 | 6.0 | 5.95 | 4.0 |
| UA | 1 | 1 | 1 | 1 |
| LA | 0 | 0 | 0 | 0 |
| $\tau_i$ (ms) | 50000 | 20000 | 20000 | 20000 |
| $\tau_\mathrm{d}$ (ms) | 50000 | 20000 | 20000 | 20000 |

| **Parameter** | **Post-synaptic (Oscillation, Anti-STDP)** | **Intracellular (Oscillation, Anti-STDP)** | **Pre-synaptic (Oscillation, Anti-STDP)** |
|---|---|---|---|
| α | 300.0 | 10.5 | 1100000.0 |
| β | 0.0041 | 0.005 | 0.25 |
| *b* | 60.0 | 400 | 80.0 |
| *c* | 0.2 | 0.2 | 0.25 |
| *d* | 0.8 | 0.7 | 0.65 |
| *e* | 15.0 | 8.0 | 80.0 |
| $\tau_r$ (ms) | 13000 | 6000 | 2500 |
| $\tau_a$ (ms) | 60000 | 100000 | 10000 |
| *w* | 70.0 | 100.0 | 100.0 |
| *I* | -20.0 | -20.0 | -18.0 |
| θ | 0 | 0 | 0 |
| ε | 0 | 0 | 0 |
| Max_rate | 10.7 | 5.95 | 5.5 |
| UA | 1 | 1 | 1 |
| LA | 0 | 0 | 0 |
| $\tau_i$ (ms) | 50000 | 20000 | 20000 |
| $\tau_\mathrm{d}$ (ms) | 50000 | 20000 | 20000 |

**Table S7. Initial values in the representative models with sleep-wake dynamics**

The initial values for channel or receptor conductance, or coefficient of pre-synaptic activation and variables of sleep-wake dynamics were as follows.

| | **Fig. 4 (Post-synaptic)** | **Fig. S12 (Intracellular)** | **Fig. S13 (Pre-synaptic)** | **Fig. S14 (bistable)** |
|---|---|---|---|---|
| conductance or coefficient (original ×) | $10^{-0.9}$ | $10^{-0.4}$ | $10^{-1.8}$ | $10^{-0.9}$ |
| ***r*** | 0.9 | 0.1 | 0.9 | 0.9 |
| ***a*** | 0.1 | 0.9 | 0.1 | 0.1 |
| $\xi$ | 0 | 0 | 0 | 0.1 |

**Table S8. The representative parameter sets for synaptic learning rules**

| **Parameter** | **Fig. 1 *B* and *C*** | **Fig. 4 (STDP) , S14 and S17*B*** | **Fig. 4 (Anti-STDP)** | **Fig. S17*A*** | **Figs. S7** and **S12 (STDP)** |
|---|---|---|---|---|---|
| $\theta_p$ | 1.16 | 1.18 | 0.67 | 1.18 | 1.2 |
| $\theta_d$ | 0.70 | 0.76 | 0.97 | 0.7 | 0.75 |
| $\gamma_p$ | 570 | 808 | 2522 | 4026 | 1324 |
| $\gamma_d$ | 161 | 222 | 7218 | 1074 | 350 |
| $\tau_{pre}$ | 36.7 | 9.83 | 3.7 | 25.8 | 7 |
| $\tau_{post}$ | 20.4 | 17.3 | 8.77 | 16.7 | 16.6 |
| $\sigma$ | 1.99 | 2.04 | 4.86 | 5.55 | 1.59 |
| $\tau_s$ | 270276 | 1033558 | 1441956 | 1553063 | 1554318 |

| **Parameter** | **Fig. S13 (STDP)** | **Fig. S12 (Anti-STDP)** | **Fig. S13 (Anti-STDP)** | **Fig. S15 (STDP)** |
|---|---|---|---|---|
| $\theta_p$ | 1.16 | 0.68 | 0.71 | 1.03 |
| $\theta_d$ | 0.74 | 1.18 | 1.01 | 0.65 |
| $\gamma_p$ | 4284 | 1580 | 2797 | 2862 |
| $\gamma_d$ | 745 | 7670 | 6816 | 926 |
| $\tau_{pre}$ | 5.77 | 2.41 | 5.55 | 49.1 |
| $\tau_{post}$ | 11 | 12.2 | 10.8 | 17.3 |
| $\sigma$ | 4.09 | 5.17 | 3.51 | 8.92 |
| $\tau_s$ | 985343 | 2254185 | 1519975 | 1073719 |

#### References


1. M. Graupner, N. Brunel, Calcium-based plasticity model explains sensitivity of synaptic changes to spike pattern, rate, and dendritic location. *Proc. Natl. Acad. Sci. U. S. A.* **109**, 3991–3996 (2012).

2. B. L. Sabatini, T. G. Oertner, K. Svoboda, The life cycle of Ca(2+) ions in dendritic spines. *Neuron* **33**, 439–452 (2002).

3. B. O. Watson, D. Levenstein, J. P. Greene, J. N. Gelinas, G. Buzsáki, Network Homeostasis and State Dynamics of Neocortical Sleep. *Neuron* **90**, 839–852 (2016).

4. Watson BO, Levenstein D, Greene JP, Gelinas JN, Buzsáki G. (2016); Multi-unit spiking activity recorded from rat frontal cortex (brain regions mPFC, OFC, ACC, and M2) during wake-sleep episode wherein at least 7 minutes of wake are followed by 20 minutes of sleep. Crcns.org.http://dx.doi.org/10.6080/k02n506q.

5. D. Levenstein, G. Buzsáki, J. Rinzel, NREM sleep in the rodent neocortex and hippocampus reflects excitable dynamics. *Nat. Commun.* **10**, 2478 (2019).

6. M. Steriade, I. Timofeev, F. Grenier, Natural waking and sleep states: a view from inside neocortical neurons. *J. Neurophysiol.* **85**, 1969–1985 (2001).

7. V. Crunelli, M. L. Lörincz, A. C. Errington, S. W. Hughes, Activity of cortical and thalamic neurons during the slow (<1 Hz) rhythm in the mouse in vivo. *Pflugers Arch.* **463**, 73–88 (2012).

8. F. Tatsuki, *et al.*, Involvement of Ca(2+)-Dependent Hyperpolarization in Sleep Duration in Mammals. *Neuron* **90**, 70–85 (2016).

9. F. Helmchen, K. Imoto, B. Sakmann, Ca2+ buffering and action potential-evoked Ca2+ signaling in dendrites of pyramidal neurons. *Biophys. J.* **70**, 1069–1081 (1996).

10. S. Song, P. J. Sjöström, M. Reigl, S. Nelson, D. B. Chklovskii, Highly nonrandom features of synaptic connectivity in local cortical circuits. *PLoS Biol.* **3**, e68 (2005).
